# Evidence-based guidelines for improving network detectability in rodent fMRI

**DOI:** 10.1101/2025.05.19.653309

**Authors:** Mila Urosevic, Gabriel Desrosiers-Grégoire, Jérémie P. Fouquet, Gabriel A. Devenyi, Daniel Gallino, Yohan Yee, M. Mallar Chakravarty

## Abstract

Mouse resting-state functional magnetic resonance imaging (rs-fMRI) is an increasingly popular tool for probing brain activity under experimental manipulations; however, there remains considerable variability in data quality throughout the field. There is a need for an accessible set of acquisition guidelines such that a baseline level of data quality can be attained regardless of domain expertise or specialized equipment (i.e. in anesthetized, free-breathing mice). In particular, there is a gap in the literature regarding the interpretation of physiological parameters as markers of anesthetic depth, and ultimately, data quality. To this end, we developed a set of acquisition guidelines after examining whether continuous physiological variables predict network detectability above and beyond categorical external variables (anesthetic dose, session, time) in C57Bl/7 and C3HeB/FeJ mice anesthetized with isoflurane-dexmedetomidine. Standard physiological metrics (respiration rate and heart rate) did not predict network detectability above and beyond anesthetic dose but instead depended strongly on strain and subject, thus we advise against tuning anesthesia based on respiration or heart rate when the goal is obtaining clear resting-state networks. The most important predictor of improved network detectability was a low isoflurane dose of 0.23%, hence we recommend that researchers prioritize piloting the minimal possible isoflurane dose for their mouse model. In summary, our work examines the contributions from sources of variability that impact rs-fMRI data quality and synthesizes the findings into practical guidelines to help experimenters adapt their acquisition protocols and improve data quality.

## Introduction

Resting-state functional magnetic resonance imaging (rs-fMRI) is rapidly becoming a key tool in preclinical neuroscience; serving as a non-invasive method of imaging brain dynamics in rodents undergoing experimental manipulations (e.g. lesions, genetic, chemogenetic or optogenetic modifications) (Desai et al. 2011; Gozzi and Schwarz 2016; Mandino et al. 2019; Giorgi et al. 2017), and as an intermediate modality facilitating the cross-species translation of findings derived from invasive techniques (e.g. Ca2+ imaging, neural recordings) (Schlegel et al. 2018; Lake et al. 2020) to humans (Pagani et al. 2023). The rising popularity of mouse rs-fMRI (Xu et al. 2022) suggests an increase in the number of non-experts seeking to apply the technique. However, the acquisition of mouse rs-fMRI data poses unique challenges beyond those in human fMRI. Amongst these is the need to keep the mouse immobile, typically achieved using either anesthesia or restraint of awake animals. The use of anesthesia is accompanied by undesirable secondary effects: alterations in patterns of brain activity (Thompson, Pan, and Keilholz 2015; Sirmpilatze et al. 2021; Gutierrez-Barragan et al. 2022), impacted neurovascular coupling (Thrane et al. 2012; Gao et al. 2017), impaired physiological regulation (Sicard and Duong 2005; Ramos-Cabrer et al. 2005; A. R. Steiner, Rousseau-Blass, and Schroeter 2021; Reimann and Niendorf 2020) and, in general, complicates comparisons to awake human studies. Meanwhile, recent awake fMRI protocols attempting to circumvent the downsides of anesthesia are often invasive (require surgical head fixation), time-intensive (4-28 days of training), may induce acute or chronic stress in the mice, do not lend themselves to high-throughput imaging studies, and do not guarantee high data quality (reviewed in (Mandino et al. 2024)). Thus, neither anesthetized nor awake imaging offer an infallible approach, and the choice between the two depends on the scientific goal of the study (Gozzi et al. 2025).

The difficulty of balancing anesthesia, motion and stress is further evidenced by the variability in data quality across mouse rs-fMRI datasets. A multi-centre study found that a significant proportion of mouse rs-fMRI scans carried important signatures of confounds (e.g. motion) or lacked the expected network connectivity (Grandjean et al. 2020). Although scans with confounded connectivity could be improved via confound correction, scans lacking network connectivity altogether were usually unfit for use even after optimized confound correction (Desrosiers-Grégoire et al. 2024). Therefore, the problems with mouse rs-fMRI cannot be solved via improvements in confound correction alone and must also be addressed at the level of acquisition. Indeed, a recent review called for an approach for identifying and achieving a consistent brain state across mouse fMRI labs (Reimann and Niendorf 2020). To this end, there is a need for accessible acquisition guidelines to improve reproducibility and reliability within the field, mitigate false positives and reduce effort spent on troubleshooting. Ideally, such guidelines would focus on the anesthetized, free-breathing approach in order to maximize accessibility to non-experts.

A key source of confusion with anesthetized, free-breathing approaches is the interpretation of physiological parameters such as respiration rate (RR) and heart rate (HR). Anesthesia impacts physiological parameters (Reimann and Niendorf 2020; Aline R. Steiner et al. 2020), hence when they are not controlled through mechanical ventilation (Shim, Lee, and Kim 2020), they vary across mice and time. It is commonly assumed that this variability in physiology may indicate differing anesthetic depths and thus multiple publications encourage monitoring mouse physiology (Constantinides, Mean, and Janssen 2011; Jonckers et al. 2015; Aline R. Steiner et al. 2020; Navarro et al. 2021) and adjusting anesthetic dose to preserve certain physiological values (Petrinovic et al. 2016; Reimann and Niendorf 2020). However, there are no specific recommendations for *how* to adjust anesthetic dose as a function of physiology and little evidence that doing so on a per mouse basis would improve reproducibility rather than reducing it.

Accordingly, the goal of this study is to produce an evidence-based set of guidelines for improving rs-fMRI network detectability in anesthetized free-breathing mice by answering the following questions: Which variables are the most important for predicting network detectability - can continuous physiological variables be more predictive than categorical external variables (anesthetic dose, session)? Can the relationship between anesthesia and network detectability be explained by physiology or motion? Based on the answers to these questions, we seek to guide researchers on whether anesthetic dose should be adjusted on a per mouse basis, as well as how to interpret and act on physiological metrics. In terms of anesthetic paradigm, we focus specifically on optimizing a combined isoflurane-(dex)medetomidine paradigm based on the extensive comparisons of anesthetic types previously performed (Williams et al. 2010; Jonckers et al. 2014; Grandjean et al. 2014; Schroeter et al. 2014; Bukhari et al. 2017; Paasonen et al. 2018). We also examine the generalizability of our findings across sexes and two background strains (C57Bl/6 and C3HeB/FeJ). Our results are synthesized into a set of practical acquisition guidelines (Appendix 1). These guidelines are intended to provide researchers with a starting point and intuition for implementing acquisition protocols, and can be combined with quality control tools from our group’s recently published software (RABIES (Desrosiers-Grégoire et al. 2024)) to validate data quality and refine protocols as necessary.

## Methods

### Experimental Overview

In order to examine the relative importance of physiological state, motion and anesthetic dose on network detectability, we deliberately acquired a highly varied dataset by systematically increasing the isoflurane dose from 0.23% to 0.5% to 1% over the course of a single uninterrupted (24 minute) scan. With this strategy we sought to capture a wide range of physiological and brain states as mice progressively transition to deeper anesthetic depths. Additionally, we varied the concentration of the dexmedetomidine infusion across 3 sessions per mouse, administering either 0.025 or 0.05 or 0.1 mg/kg/h in a randomized order (so that the effect of dexmedetomidine dose is not confounded by the effect of session). Thus, every mouse experienced 9 combinations of isoflurane-dexmedetomidine doses (Figure 1a). During fMRI acquisition, we recorded respiration, plethysmography and oxygen saturation, all of whom were allowed to vary freely - only body temperature was controlled at 36.5℃. Note, it is possible to also control respiration (via mechanical ventilation) and potentially also oxygen saturation (via the ratio of inspired oxygen to air) but we chose not to in case the natural values provide useful information.

**Figure 1.**
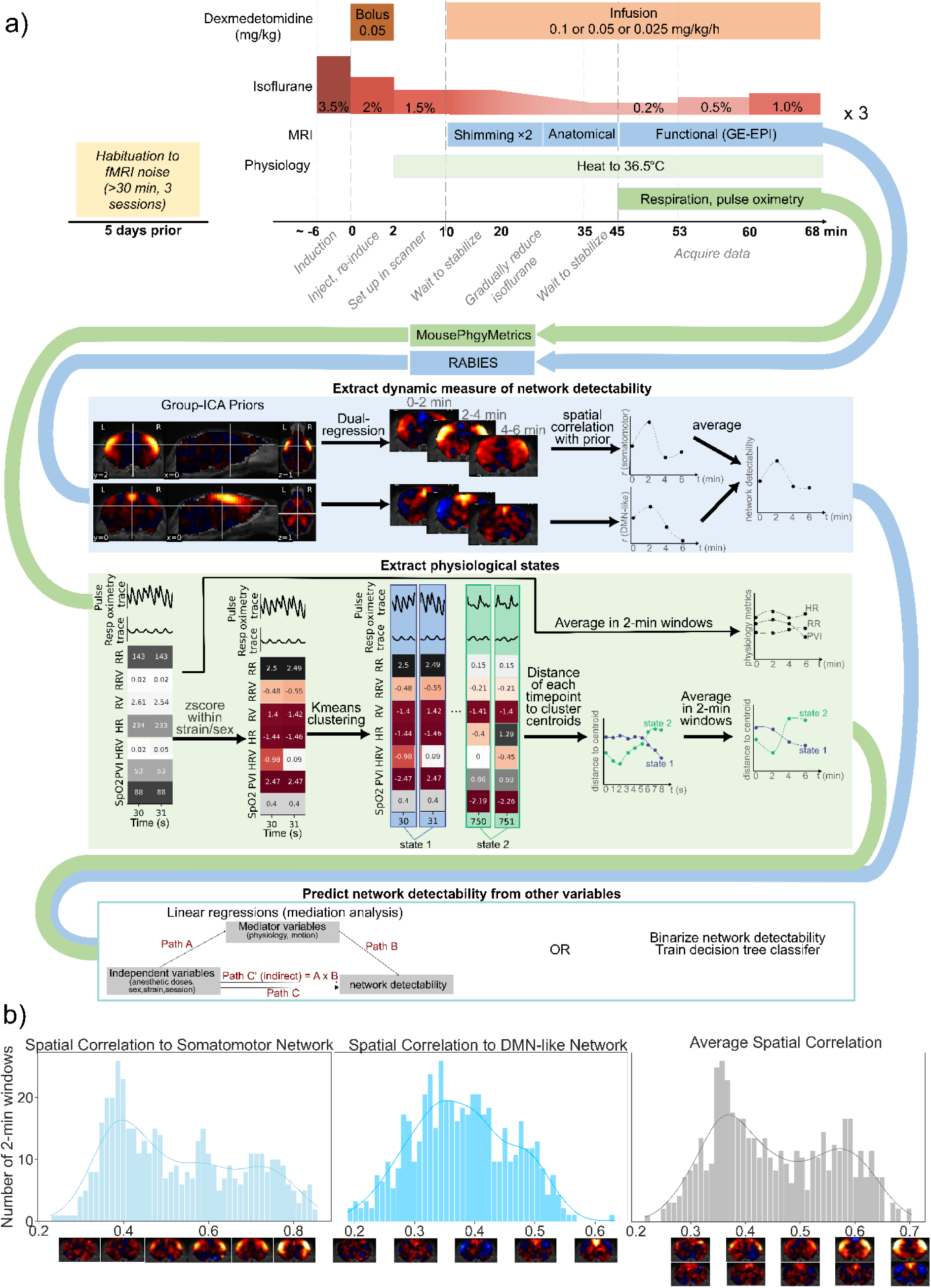
Overview of methodology and rationale. a) The methodology consisted of: acquiring simultaneous fMRI and physiology data across a range of isoflurane and dexmedetomidine doses, preprocessing fMRI data with RABIES and physiological data with our custom software (MousePhgyMetrics), extracting a measure of network detectability from the fMRI data in 2-min windows, extracting physiological metrics as well as physiological states (co-occuring patterns of physiology) also within 2-min windows, then predicting network detectability from the other acquisition-related variables. The predictions can be done using linear regressions in a mediation analysis framework, to parse the direct impact of anesthesia and demographics from the indirect impacts via physiology/motion. Alternatively, network detectability can be predicted using decision trees to additionally examine non-linear dependencies between variables. Acronyms for physiological variables are defined in Table 1. b) Histograms displaying the distribution of network detectability values (for the somatomotor and DMN-like networks, as well as their average) across all time windows and subjects. The dataset was highly varied, containing a range of network detectability values for the somatomotor and DMN-like networks, as well as their average. Spatial maps along the x-axis are representative examples of the fitted network representations for each corresponding correlation value.

**Table 1:** Physiological metrics that were calculated with the MousePhgyMetrics toolkit from the respiratory (dark green) and pulse oximetry (light green) traces. All of the metrics were chosen due to previously published evidence that they are markers of anesthetic depth or proxies for more fundamental (yet more invasive) measures.

| Physiological metric | Relevance | Calculation |
| --- | --- | --- |
| Respiration rate (RR) | Usually decreases proportionally to anesthetic dose due to respiratory depression. Also reflects blood levels of oxygen and carbon dioxide. | Number of peaks in a 2 s rolling window, divided by window length. |
| Heart rate (HR) | Reflects blood pressure and blood levels of oxygen and carbon dioxide. May either increase (under isoflurane) or decrease (under medetomidine) in response to anesthesia depending on the vasodilatory properties of the anesthetic. |  |
| Respiration rate variability (RRV) | Increases are indicative of elevated blood pressure (Anderson et al. 2008) and poor prognosis in patients (Garrido et al. 2018). | Standard deviation of the time between peaks in a rolling window of 4 peaks. |
| Heart rate variability (HRV) | Reductions indicate autonomic nervous system dysfunction and/or deep anesthesia (Sakuma, Ueda, and Koide 1985; Mazzeo et al. 2011). |  |
| Respiratory variation (RV) | Proxy for inspired volume over time (Chang, Cunningham, and Glover 2009). Similar measures have been shown to drive network-like neural activity (Tu and Zhang 2022). | Standard deviation of the respiratory waveform in a rolling window of 3 s (Chang, Cunningham, and Glover 2009). |
| Plethysmography variability index (PVI) | Relates to blood volume (Shamir et al. 1999), intrathoracic pressure, stroke volume and cardiac output (Masimo Corporation 2013). | Percent change between the maximum and minimum heart beat amplitudes, in a rolling window of 9 heart beats (Sano et al. 2018). |

From the preprocessed fMRI data, we computed a dynamic measure of network detectability by first partitioning data into 2-minute time windows then quantifying how well-represented two canonical networks (somatomotor and DMN-like) are in each window via a spatial correlation to the original “ground-truth” maps (Figure 1a). Our measure of network detectability was the average spatial correlation of the two networks. As intended, network detectability varied highly across the dataset, with certain time windows almost perfectly recapitulating both networks, while other windows seemingly contained only noise (Figure 1b). To understand which factors distinguish the high network detectability windows from the poor ones, we sought to predict network detectability from isoflurane dose, dexmedetomidine dose, strain, sex, subject, session (correlated with age), motion levels, physiological variables as well as physiological states (co-occuring patterns of physiological variables). This was achieved using two complementary approaches: mediation analysis (consisting of three linear mixed effects regressions) and a decision tree.

### Verification of the scanner’s temporal stability

Prior to examining how *in vivo* factors impact network detectability, we first verified the temporal stability of our Bruker Biospec 7T preclinical scanner (70/30 USR; 30-cm inner bore diameter, located at the Douglas Research Centre’s Cerebral Imaging Centre in Montreal, QC, Canada) to rule out significant temporal noise or instabilities that would confound interpretation of downstream results. Such verifications are recommended in the human fMRI literature (Friedman and Glover 2006; Kayvanrad et al. 2021), but have thus far never been reported for preclinical scanners. To this end, we assembled an agarose phantom in a 3mL syringe (2% agarose, 0.9% sodium chloride, 0.1% sodium azide, 2.0× 10^−5^% manganese chloride) (Glover 2003; Friedman and Glover 2006) such that the relaxation properties approximate those of the mouse brain. Temporal fluctuations in the phantom’s fMRI timeseries will be attributable to temporal noise and biases in the scanner electronics.

Scanning was performed with a surface cryogenic transmit/receive coil (CryoProbe™; Bruker, Ettlingen, Germany). B0 mapping and shimming were performed with the MapShim and FieldMap functions in Paravision 6.0.1 using a cylindrical shim volume. fMRI scanning was performed with a gradient echo-echo planar imaging (GE-EPI) sequence and the following parameters: TE = 15 ms, TR = 1000 ms, bandwidth = 300480.7692 Hz, matrix = 75×40, FOV = 18.75×10 mm, anti-aliasing = 1.28×1, 26 interlaced slices, slice thickness = 0.5 mm, 360 repetitions. The phantom was scanned four times, with sessions spaced 1 month apart.

The data was examined for temporal instabilities using a custom toolbox (https://github.com/CoBrALab/fMRI_phantom_analysis). The toolbox performed a principal component analysis (PCA) decomposition along the temporal dimension in order to extract temporal patterns that explain the most temporal variance in the data, as well as their corresponding spatial maps. Additionally, the toolbox computed standard stability metrics that are used in the human fMRI literature, including temporal signal to noise ratio (tSNR), drift and percent fluctuation (the ratio of the standard deviation of fluctuations around the mean, to the mean of the timecourse) (Friedman and Glover 2006). Overall, we identified a periodic temporal instability with a structured spatial pattern that was pinpointed to the IECO gradient amplifiers, as well as a slowly varying cubic instability right under the CryoProbe™ (Fig. S1). The remaining temporal variance was explained by seemingly random noise. Overall, the temporal instabilities and noise were well below the expected percent change in BOLD signal (0.20±0.04% vs 1-2.5% change in BOLD (Desai et al. 2011; Jung, Shim, and Kim 2019)) and should not significantly corrupt *in-vivo* BOLD recordings. As such we deemed the scanner to be stable for downstream analyses of network detectability.

### Animals

Data was acquired on 16 adult mice (8-9 weeks old at the first session) from two common background strains: C57Bl/6J and C3HeB/FeJ (8 mice per strain, 4 male/4 female), sourced from the Jackson laboratory. Animals were housed 4 per cage at the Douglas Research Centre’s animal facility (Montreal, QC, Canada) under standard conditions (12-hour light/dark cycle) with *ad libitum* access to food and water. All animal experiments were carried out in accordance with the Canadian Council on Animal Care and approved by the McGill University Animal Care Committee (Montreal, QC, Canada).

### Mouse preparation and anesthesia

Mice were habituated to scanner sounds 5 days prior to the first scanning session. Habituation involved placing the mouse cages adjacent to the scanner room for 30+ minutes/session over 3 days while the scanner was in use for other fMRI experiments. The decision to perform sound habituation was made out of concern that mice may wake due to stress under the very light anesthesia (as suggested in (Nasrallah, Tay, and Chuang 2014)).

On the scanning day, anesthesia was induced at 3.5% isoflurane in 100% oxygen, a dexmedetomidine (Dexdomitor, Orion Corporation, Espoo, Finland) bolus of 0.05 mg/kg was injected intraperitoneally and ophthalmic ointment was applied. Mice were transferred to the scanner room where the needle for continuous dexmedetomidine infusion was inserted intraperitoneally. The mice were secured within the Bruker nose cone via a bite bar. 1.5% isoflurane was delivered in a 80/20% air/oxygen mixture at a flow rate of ∼1 L/min. Mice were warmed to 36.5℃ using an air heater at 40% power receiving feedback on body temperature from a rectal temperature probe (SA Instruments Inc., NY, USA). Respiration and pulse oximetry were monitored via a pneumatic sensor and an ankle pulse oximeter, respectively (SA Instruments Inc., NY, USA). Silicone ear plugs (Mack’s, MI, USA) were inserted, and a thin layer of padding was secured on the surface of the head in order to reduce room for movement. The continuous dexmedetomidine infusion (0.025, 0.05 or 0.1 mg/kg/h) was started 10 minutes following the initial bolus. The isoflurane level was reduced stepwise during anatomical scanning until it reached 0.23%. The fMRI scan was initiated after 10 minutes at 0.23% and at least 45 minutes post-bolus (Pradier et al. 2021).

Isoflurane was increased stepwise from 0.23% to 0.5% to 1% every 8 minutes throughout the uninterrupted fMRI scan. The same anesthesia regime was maintained regardless of the mouse’s respiration and heart rate. Only in the event of clear and repeated signs of motion (visible as large spikes or voltage shifts in the real-time respiration trace that do not appear to have a physiological origin) indicating that the mouse might be awakening was the isoflurane at 0.23% skipped. After the scan, mice were injected subcutaneously with atipamezole hydrochloride (Antisedan, Orion Corporation, Espoo, Finland) to facilitate recovery from dexmedetomidine (the dose of atipamezole was equal to 10 times the total dose of dexmedetomidine administered in milligrams). The same process was repeated for 3 sessions using a different dexmedetomidine concentration each time, with a 3-10 day gap between consecutive sessions. See Figure 1a.

### Image Acquisition

Scanning was performed with the same equipment as during the phantom experiments for the verification of scanner stability. Two rounds of B0 mapping and shimming were performed using an ellipsoidal shim volume (covering as much of the brain as possible without entering the skull or olfactory bulbs) where each round was followed by a point-resolved spectroscopy (PRESS) acquisition (TE = 16.5 ms, TR = 2500 ms, number of averages = 10, excitation FA = 90°, refocusing FAs = 180°, voxel size = ∼ 634 mm (depending on the shim volume), number of sampling points = 2048, nucleus = 1H, scan time = 25 s) to confirm that homogeneity improved and that the linewidth’s full-width at half-maximum (FWHM) was ∼25 Hz or less. An anatomical T2-weighted image was acquired with a rapid acquisition with relaxation enhancement (RARE) sequence (TE = 22.4109 ms, RARE factor = 4, TR = 3500 ms, number of averages = 2, 1 echo image, excitation FA = 90°, refocusing FA = 180°, with flip back, matrix = 187×100, FOV = 18.75×10 mm, 39 slices, voxel size = 0.1×0.1×0.33 mm, scan time = 5:22 minutes). fMRI scanning was performed with the same sequence used in the phantom experiment, extended to 1440 repetitions.

### fMRI Preprocessing

All fMRI preprocessing was conducted with the open-source RABIES software (version 0.5.0) (Desrosiers-Grégoire et al. 2024). The anatomical T2w images underwent: non-local means denoising (Manjón et al. 2010), iterative inhomogeneity correction with the N4ITK algorithm (Sled, Zijdenbos, and Evans 1998; Tustison et al. 2010), cross-subject alignment via iterative non-linear registration to the dataset consensus average (Avants et al. 2011; Germann et al. 2025), followed by non-linear registration of the consensus average to the Dorr-Steadman-Ullmann-Richards-Qiu-Egan (DSURQE) atlas (Dorr et al. 2008; Steadman et al. 2014; Richards et al. 2011; Ullmann et al. 2013). The functional EPIs underwent a preliminary motion realignment, derivation of a volumetric EPI from the trimmed mean across EPI volumes, denoising of the volumetric EPI with non-local means algorithm (Manjón et al. 2010), estimation of head motion parameters from the realignment of EPI volumes to the volumetric EPI via rigid registration (Avants, Tustison, and Song 2009), inhomogeneity correction with the N4ITK algorithm (Sled, Zijdenbos, and Evans 1998; Tustison et al. 2010), distortion correction via non-linear registration to the corresponding anatomical scan (S. Wang et al. 2017).

Preprocessed timeseries are generated in atlas space (i.e. common-space) by concatenating previously computed transforms (i.e. motion realignment, distortion correction, resampling onto dataset consensus average, and resampling onto atlas space) and resampling each original EPI volume with a single interpolation step (Esteban et al. 2019).

Confound correction was executed using the RABIES software (Desrosiers-Grégoire et al. 2024) on the common-space fMRI time-series. First, high motion timepoints were censored (using a threshold of FD > 0.05 mm, and DVARS > 2.5 standard deviations), followed by voxel-wise linear detrending, bandpass filtering (0.01-0.3 Hz), and removal of edge artifacts by discarding 30 s from either end of the time-series (Power et al. 2014). To combine filtering and censoring, missing frames are simulated while preserving the frequency composition of timeseries and removed again post-filtering as described in (Power et al. 2014). Nuisance signals were filtered likewise to prevent re-introduction of previously filtered artefacts (Lindquist et al. 2019), and then regressed, and consisted of: 6 rigid motion parameters, mean white matter signal, mean cerebrospinal fluid signal, mean vascular signal. We opted for a minimal confound regression scheme to decrease the likelihood of removing relevant network information (Bright and Murphy 2015). Finally, a spatial Gaussian smoothing kernel (Abraham et al. 2014) was applied at 0.5 mm FWHM (as in (Gutierrez-Barragan et al. 2022)). The suitability of this confound correction scheme was evaluated by examining the spatial and temporal diagnosis figures generated by the ‘analysis’ function in RABIES (Desrosiers-Grégoire et al. 2024). Finally, sections of the scan that were excluded from physiology analysis due to noisy physiological recordings were also censored from the fMRI timeseries.

Identical preprocessing and confound correction steps were also applied to an external, high-quality dataset from (Grandjean and Yeow 2020) that will serve as a reference for ‘ground-truth’ networks. The only additional processing step for this dataset was to resample the fMRI data with RABIES to 0.25×0.25×0.5 mm to match the resolution of our local dataset.

### Calculation of network detectability

We focused on the somatomotor (also referred to as the lateral cortical network (Gozzi and Schwarz 2016)) and DMN-like networks because they are the two most discussed networks in the literature, making them a useful minimal benchmark for rs-fMRI data quality (Desrosiers-Grégoire et al. 2024). ‘Ground truth’ spatial maps (spatial priors) of the networks were obtained from a group independent component analysis (ICA) decomposition (Beckmann and Smith 2004) on an external fMRI dataset acquired on 9 awake and 10 anesthetized female adult C57Bl/6 mice (Grandjean and Yeow 2020), chosen because for its previously reported high quality (Desrosiers-Grégoire et al. 2024). The priors were derived on an external dataset because deriving them on the present dataset with its intentionally heterogeneous quality may produce biased or slightly corrupted maps, even on a group level. The spatial priors were smoothed with a Gaussian filter (FWHM = 0.5 mm) prior to connectivity measurements.

The spatial priors were fitted to each scan (within 2-minute windows) via dual-regression, as implemented in RABIES (Desrosiers-Grégoire et al. 2024). The window length was chosen from calcium-imaging evidence that sufficient neuronal coactivations should occur sometime within any 2 minutes (Matsui, Murakami, and Ohki 2016). To quantify network detectability, we calculated the Pearson correlation coefficient between the original spatial priors and their fitted representations in each window (smoothed with a 0.3 mm FWHM Gaussian kernel), then averaged the spatial correlations over the two networks. Thus, the network detectability metric indicates to what extent either the somatomotor or DMN-like networks were active in a given window.

### Physiological Data Acquisition

In order to examine whether network detectability can be predicted from physiological metrics, we acquired respiratory and pulse oximetry recordings during the fMRI scan. The raw respiratory and pulse oximetry traces were captured at 225 and 450 samples/second respectively with the signal breakout module and WaveGrab software (SA instruments Inc., NY, USA). Oxygen saturation was exported at 1 sample per second using TrendMap (SA instruments Inc., NY, USA). All data is openly available at: https://doi.org/10.5281/zenodo.12637274.

### Physiological data processing and metric extraction

Physiological data preprocessing and metric calculation was performed with the MousePhgyMetrics toolbox (Python 3.8.8), which was developed explicitly for the present study but is now open and fully documented online (https://github.com/CoBrALab/MousePhgyMetrics). This custom toolkit was developed as we found existing softwares did not output the range of metrics that we desired or had defaults that were not well-suited to quantifying mouse physiology. Using MousePhgyMetrics, we first smoothed the physiological traces in a 90 ms Gaussian rolling window and detrended the respiratory trace by subtracting the mean in a 1s window (Kalthoff et al. 2011; Noto et al. 2018).

Subsequently, we computed the wavelet transform of the physiological trace, obtaining a data-driven and noise-robust overview of the respiration/heart rate throughout the acquisition. Next, breaths/heart beats were extracted from the physiological traces using default peak detection parameters (Scipy signal package (Virtanen et al. 2020)) - these defaults had been carefully selected during preliminary analyses for being suitable for the majority of acquisitions. We performed a visual quality control (QC) of the outputs, identified unusual acquisitions for which the default parameters did not accurately extract peaks via comparison with the wavelet transform, and modified the parameters accordingly until the visual QC was satisfactory. We were careful to properly include ectopic beats in the heartbeat detection. Where the trace was too noisy and it was hard to visually identify the correct breaths/heart beats, the data was excluded (see Table S1 for a complete list of excluded data). Finally, from the extracted peaks, we computed a set of physiological metrics that are known to be useful indicators of physiological state and/or anesthetic depth (Table 1) at each timepoint. The SpO_2_ values underwent minor preprocessing to remove the bottom 1% and top 99% outlier values and interpolate missing values. The metrics were averaged within 2-minute time windows.

### Extracting physiological states via clustering

The individual physiological metrics, while interpretable, may have limited predictive power. Similarly to how brain activity is known to transition through different states over time, it is plausible that the body also transitions through a set of physiological states (i.e. co-occuring patterns of respiration and pulse oximetry), particularly as the mouse experiences different anesthetic depths. Thus, a physiological state (e.g. high RR combined with low HR) may be more indicative of anesthetic depth and network detectability than any single physiological metric. To examine this possibility, we extracted physiological states by performing K-Means clustering of physiological metrics on a timepoint level, borrowing from standard co-activation pattern (CAP) analysis for brain states (Liu et al. 2018).

K-Means clustering was performed in Python 3.8.8 using a custom function where the distance metric is Pearson correlation across features (to mimic CAP analysis (Bolton et al. 2020)). The input physiological metrics were z-scored within each subgroup of strain and sex to account for the differences in physiology between strains/sexes. K-Means clustering initializes *n* random cluster centroids, assigns each point to the cluster that it is most correlated with, updates centroids as the mean of assigned points, and iterates until convergence. The optimal number of clusters was determined to be 5 from a comparison of silhouette plots, which indicate how close each point is to its assigned cluster compared to neighboring clusters (Rousseeuw 1987). We extracted the correlation of each timepoint to each of the final 5 cluster centroids and averaged these correlations in 2-minute windows.

### Predicting network detectability with linear mediation analysis

Our goal was to examine whether network detectability in a given window is best predicted by continuous physiological variables or categorical external variables (demographics, anesthetic doses, session, time). However, the physiological variables will themselves be influenced by the external variables, thus we must test whether they are predictive *above and beyond* the external variables.

This is best achieved with a mediation analysis (Baron and Kenny 1986). The first step is to confirm that the physiological variables are influenced by the external variables via linear regression (path A, equations 1-7). Next, the predictive role of the physiological variables is determined by comparing two regression models: a model where network detectability is predicted only from external variables (Path C, equation 8) and one where it is predicted from both external (path C’) and physiological (path B) variables (equation 9). If the effect of a given external variable decreases or disappears when the physiological variables are added (C’-C), it means that certain physiological variables are mediators - they partially or totally explain the impact of the independent variables on network detectability. These mediators not only explain the relationship, they are actually better predictors than the external variables because they predict network detectability even when controlling for the external variables (significant path B), i.e. they predict network detectability even within a single condition of each external variable. Thus, with this approach we can determine the added value of monitoring physiology. Note that in our models, we also controlled for the random effect of subject and covaried for possible interactions between demographics and anesthetic dosage. In total, the mediation analysis consists of the following 9 linear models:

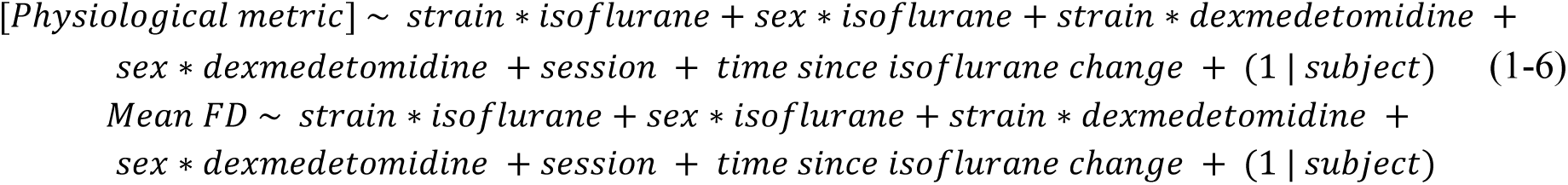

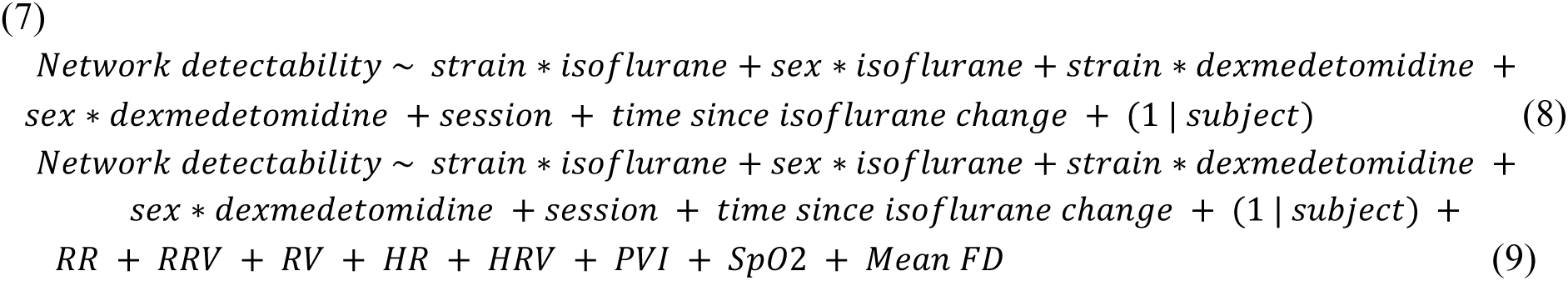

The analysis was performed in R 4.2.0. The ordered categorical variables (i.e. isoflurane dose, dexmedetomidine dose and session order) were coded as backward difference contrasts (Venables and Ripley 2003), such that each level was compared to the previous level (“R Library Contrast Coding Systems for Categorical Variables,” n.d.). The network detectability metric underwent a Fisher z-transformation (Revelle 2022) to reduce the skewness and a log transformation to further approximate a normal distribution. All other continuous variables were standardized to a mean of 0 and standard deviation of 1. Weight and age were excluded due to collinearity with other demographics (Lüdecke et al. 2021). The linear regressions were modeled using the *brms* package for Bayesian regression modeling with Stan (Bürkner 2017, 2018; Clark 2019; Bürkner 2021). It was necessary to use Bayesian statistics because standard packages were not flexible enough for the desired model. The Bayesian approach produced coefficient estimates as well as confidence intervals; if the confidence interval for a given result did not cross zero, then the null hypothesis could be rejected, indicating a significant result. As such, the approach did not produce p-values and did not require correction for multiple comparisons (G. D. Garcia 2024), moreover all 9 regressions could be performed simultaneously. We evaluated the relative importance of each predictor by comparing the increase in variance explained (*R^2^*) obtained by adding the predictor to the model. Since additional *R^2^* depends on which order the variables were added in, the average across all orderings (i.e. the LMG metric, named after its authors (Lindeman, Merenda, and Gold 1980)) was computed with the *relaimpo* package (Groemping 2007).

### Predicting network detectability with decision trees

To provide researchers with an intuitive understanding of the consequences of their acquisition decisions, we also implemented a decision tree classifier. Decision trees are constructed layer-by-layer by splitting the dataset based on the features that will yield the purest subsets that best distinguish windows containing a recognizable network from those that do not.

The decision tree classifier was implemented in Python 3.11.7 with scikit-learn 1.3.2 (Pedregosa et al. 2011). First, because decision is a binary classification problem, the network detectability metric was binarized into: recognizable network present (network detectability ≥ 0.45) and no recognizable network (network detectability <0.45), where the threshold was determined from the trough of the bimodal distribution (Figure 1b). The dataset underwent a stratified split into training and test sets (75%/25% split), where all windows from a given scan were assigned to the same set in order to avoid leaking information into the test set.

Physiological metrics that were found to depend strongly on strain or sex, were z-scored within each strain/sex combination - this was done so that the decision tree would not simply pick out a strain or sex by splitting on a physiological metric but would identify the usefulness of a physiological metric within a strain/sex combination (see Figs.S2 for the original means and standard deviations). The parameters for the z-scoring were determined from the train set only, then applied to both train and test, again in order to avoid leaking information into the test set. We trained decision trees across a range of hyperparameters (depth: 2-4, maximum number of leaf nodes: none or double the depth, splitter: ‘best’ or ‘random’), while the minimum number of samples per leaf was fixed at 20. The final decision tree, chosen for its balance of interpretability, generalizability and low rate of false positives, had a depth of 4, maximum of 8 leaf nodes and was constructed with the random splitter. We also computed the contribution of each feature to the decision tree by finding the mean decrease in Gini impurity obtained by splitting on that feature, averaged across all layers.

All code for the analysis that was not performed with one of the previously described toolboxes is available here: https://github.com/CoBrALab/mousefmri_acq_publication/tree/main.

## Results

### Physiology and motion metrics are primarily dependent on strain

To examine whether physiology and/or motion predict network detectability above and beyond the external variables (anesthetic doses, demographics, session, time), we began by confirming that physiological variables reflect the external variables (path A of mediation analysis, Figure 2a)

**Figure 2:**
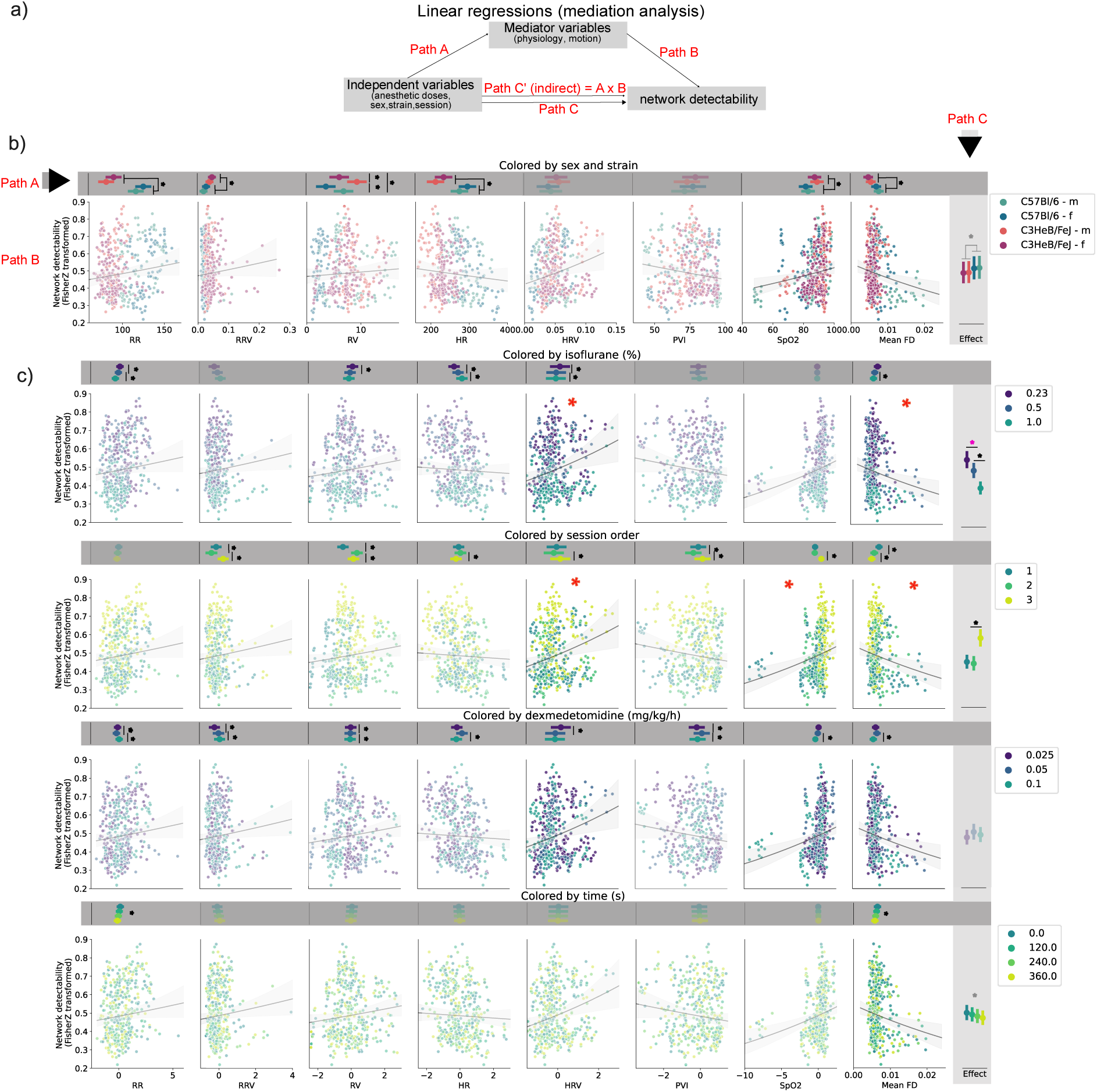
The effect of significant independent variables (strain/sex, isoflurane, session and dexmedetomidine) on network detectability, both directly and indirectly via their influence on mediators (physiology and motion). a) Diagram of mediation analysis explaining the path names, where paths A, B and C are modeled with linear mixed effects regressions, and the strength of path C’ is derived by multiplying regression coefficients from paths A and B. b) Mediation analysis results for the effects of strain and sex via all mediators. The horizontal gray band depicts the regression coefficients and 95% confidence intervals for the effects of strain/sex on each mediator (Path A). The vertical gray band along the right depicts the total effect of strain/sex on network detectability (Path C). Pointplots are faded out if there are no significant effects. The scatterplots depict the effect of each mediator on network detectability, where each point corresponds to a 2-minute window (Path B), colored by strain/sex. The mediators that are not significant (i.e. no indirect effect through path C’) are faded out. The x-axis is shown in the original scale of each variable. The star indicates significance (confidence interval doesn’t contain zero); Bayesian statistics do not produce p-values so it isn’t possible to show degrees of significance. c) Mediation analysis results for remaining independent variables. The formatting is similar to (b), except the physiological variables are now z- scored within each strain/sex subgroup, in order to highlight the effect of each independent variable regardless of strain/sex. Black significance stars indicate effects that are stable regardless of the presence of mediating variables in the model. Red significance stars indicate mediator variables that explain the relationship between the corresponding external/independent variables and network detectability. Pink significance stars indicate an effect that completely disappears when the mediators are added to the model. Gray significance stars indicate an effect that appears only after the addition of mediators to the model.

A considerable portion of the variance in physiological and motion metrics were explained by the external variables. The metrics with the highest variance explained were RR (83%), RV (82%) and HR (81%) whereas the one with the least variance explained was RRV (39%) (Table 2). Out of this total variance explained, the majority was explained by demographics — namely strain, sex and subject. Not only did strain explain a high proportion of the variance, but it also had a large and significant effect on most physiological metrics (Figure 2b).

**Table 2:** Importance of regressors in predicting physiological metrics and motion.

| % of variance explained by each regressor |  |  |  |  |  |  |  |  |  |
| --- | --- | --- | --- | --- | --- | --- | --- | --- | --- |
| X | Regressors | LMG |  |  |  |  |  |  |  |
|  |  | RR | RRV | RV | HR | HRV | PVI | SpO2 | mean FD |
| 1 | strain | 54.0 | 13.0 | 9.6 | 50.7 | 5.8 | 0.1 | 9.8 | 15.6 |
| 2 | sex | 2.1 | 2.6 | 16.1 | 4.9 | 0.7 | 3.3 | 1.3 | 0.6 |
| 3 | session | 1.3 | 3.0 | 14.6 | 4.0 | 2.8 | 1.2 | 16.9 | 8.0 |
| 4 | isoflurane | 1.6 | 0.9 | 2.2 | 3.9 | 0.8 | 0.2 | 0.2 | 0.9 |
| 5 | dexmedetomidine | 1.1 | 1.0 | 0.5 | 0.6 | 2.1 | 2.8 | 7.1 | 1.0 |
| 6 | time after isoflurane change | 0.8 | 0.2 | 0.1 | 0.1 | 0.0 | 0.1 | 0.0 | 0.5 |
| 7 | strain:isoflurane | 0.4 | 0.8 | 0.1 | 1.8 | 1.9 | 0.5 | 0.1 | 1.7 |
| 8 | sex:isoflurane | 1.6 | 0.3 | 0.1 | 0.0 | 0.1 | 0.3 | 0.1 | 0.1 |
| 9 | strain:dexmedetomidine | 1.7 | 2.0 | 12.4 | 0.2 | 1.2 | 11.1 | 4.3 | 3.1 |
| 10 | sex:dexmedetomidine | 4.4 | 3.6 | 5.7 | 0.1 | 13.0 | 8.2 | 6.2 | 1.7 |
| 11 | subject | 16.0 | 11.0 | 24.0 | 22.0 | 55.0 | 47.0 | 16.0 | 18.0 |
| 12 | Total | 83.0 | 39.0 | 82.0 | 81.0 | 76.0 | 63.0 | 58.0 | 50.0 |

The variance explained ( $R^2$ ) by a regressor is calculated as: the increase in model $R^2$ when that regressor is added to the model. This value depends on the order in which that regressor is added to the model, thus the LMG metric represents the average $R^2$ across all orderings. Values were computed with the relaimpo package (Groemping, 2007). LMG values are not available for the random effect of subject, thus the $R^2$ for subject was obtained by subtracting the fixed effects $R^2$ from the total effects $R^2$ . The most important regressor for each metric is highlighted in yellow.

In particular, RR and HR were the most impacted by strain, which accounted for 50.7-54% of the variance explained (as quantified by the LMG metric i.e. the average R^2^ across all possible orders that the variable is added to the linear model). The difference in RR and HR between the two strains was easily visible even in the raw data (Fig 2b scatterplots), with the C3HeB/FeJ strain exhibiting a dramatically lower RR (β_S_=-1.573, β=-35 bpm) and HR (β_S_=-1.399, β=-62 bpm) compared to the C57Bl/6 strain (Fig 2b, path A). Other physiological metrics that depended on strain included RRV (β_S_=0.705, β=1.82× 10^−3^, LMG=13%), RV (β_S_=0.644, β=2.28× 10^−3^, LMG=9.6%), SpO_2_ (β_S_=0.618, β=6%, LMG=9.8%) and mean FD (β_S_=-0.762, β=-1.87× 10^−3^mm, LMG=15.6%). RV was additionally dependent on sex, where females had lower RV compared to males (β=-0.966, LMG=16%). The only metrics that were not dependent on either strain or sex were HRV and PVI, yet both of those showed high interindividual variability, where the random effect of subject explained a large portion of the variance (LMG=55%, LMG=47% respectively). Thus, every single physiological and motion metric varied significantly between strains, sexes or subjects.

Nevertheless, when controlling for demographics, all physiology and motion metrics also reflected anesthetic dose. A higher isoflurane dose results in slower and shallower breaths (β of-5 to - 13 bpm), faster and more regularly spaced heart beats (β of-17 to-23 bpm), and less motion (β of - 9× 10^−3^mm). The changes in RR and HR were approximately doubled when going from 0.5% to 1% isoflurane compared to going from 0.23% to 0.5%. On the other hand, increasing dexmedetomidine dose resulted in slower and more irregular heart beats (β of-29 bpm), decreased oxygen saturation (β of-10%), decreased motion (β of-1× 10^−3^mm) and conflicting effects on respiration rate (depending on the dose). See Table S1 for all the standardized effect sizes and confidence intervals. Overall, anesthetic dose explained an additional ∼10% of variance after accounting for all other variables (i.e. when added last to the model) despite its low LMG (Table 2).

Additionally, all physiology and motion metrics (except RR) were significantly impacted by session order, a variable indicating whether it was the mouse’s 1st, 2nd or 3rd time being scanned. However, the effect sizes were mild and the direction of the effect frequently reversed when going from session 2 to 3 compared to when going from 1 to 2, making it difficult to conclude how session overall impacts physiology. The effects were most robust for RV and SpO_2_, where session explained 14.6% and 16.9% of the variance respectively.

### Network detectability is primarily influenced by isoflurane dose

Next, we investigated the impact of external variables (anesthetic doses, demographics, session, time) alone on network detectability; this regression is known as path C in the mediation analysis framework (Fig 2a). In total, we were able to explain 61% of the variance in network detectability with the independent variables and their interactions. Of these, isoflurane dose was the most important predictor, with by far the largest LMG value of 18.9%. In addition to explaining the most variance, changing isoflurane dose also had the largest effect size on network detectability; increasing isoflurane from 0.23% to 0.5% lowered network detectability by β=0.08 and increasing isoflurane again from 0.5% to 1% lowered it by another β=0.23 (Fig 2c, light gray column), enough to go from a clean network pattern to an undetectable one (see Figure 1c for interpretation of network detectability values). The second most important predictor of network detectability was session order (LMG = 9.9%). There was no significant difference between the 1st and 2nd scanning sessions, but on the 3rd session, network detectability increased by β=0.27 compared to the previous session (Fig 2c, light gray column).

**Table 3:** mportance of regressors in predicting network detectability.

% of variance in network detectability explained by each regressor, computed in 3 ways
| Regressors | LMG | First | Last |
| --- | --- | --- | --- |
| independent variables |  |  |  |
| strain | 1.2 | 0.0 | 0.1 |
| sex | 0.4 | 0.9 | 0.1 |
| session | 9.9 | 16.4 | 5.2 |
| isoflurane | 18.9 | 23.3 | 15.1 |
| dexmedetomidine | 0.8 | 1.3 | 0.9 |
| time after isoflurane change | 0.4 | 0.4 | 0.4 |
| instantaneous variables |  |  |  |
| RR | 4.1 | 2.7 | 1.7 |
| RRV | 0.3 | 0.1 | 0.3 |
| RV | 0.4 | 0.2 | 0.0 |
| HR | 2.6 | 5.1 | 0.1 |
| HRV | 1.1 | 1.4 | 0.4 |
| PVI | 0.4 | 0.4 | 0.0 |
| SpO2 | 5.3 | 11.1 | 1.3 |
| mean FD | 1.5 | 2.4 | 0.8 |
| total instantaneous | 14.7 | 26.0 | 9.0 |

The variance explained ( $R^2$ ) by a regressor is calculated as: the increase in model $R^2$ when that regressor is added to the model. This value depends on the order in which that regressor is added to the model, thus we present 3 approaches. LMG is the average $R^2$ across all orderings. 'First' is when that regressor is added first. 'Last' is when that regressor is added last. Values were computed with the relaimpo package (Groemping, 2007). Interaction terms and random effects are not shown as their 'first' and 'last' contribution cannot be calculated with the package. The 'total instantaneous' row shows the additional variance explained if all the instantaneous variables are added at once, either in first or last order - it is not necessarily equal to the sum of their first and last variance explained.

### HRV, SpO_2_ and mean FD are predictive of network detectability above and beyond isoflurane dose

Finally, we tested whether any physiological variables explain the relationship between the external variables and network detectability by adding physiological variables to the path C model (path C’). Note that combining all physiological and external variables as predictors in one model also indicates which physiological variables are significant when controlling for the external variables (path B).

Indeed, the significant difference in network detectability between isoflurane doses of 0.23% and 0.5% disappeared; completely explained by HRV and mean FD. Therefore, not only did HRV and mean FD reflect changes in isoflurane dose, but they were also predictive of network detectability within a given isoflurane dose (since this model is controlling for all external variables), explaining network detectability above and beyond the effect of isoflurane. Similarly, the significance of the difference in network detectability between sessions 2 and 3 decreased from β=0.27 to β=0.20; being partially explained by HRV (β=0.06), mean FD (β=-0.04) and SpO_2_ (β=0.05). Hence, an increased HRV, decreased mean FD and increased SpO_2_ may be useful live indicators of good network detectability with the former two being potential markers of anesthetic depth.

Nevertheless, even after adding the physiological variables to the model, a strong significant difference between isoflurane of 0.5% and 1% still persisted (β=-0.186, LMG=18.9). Thus, at high doses of isoflurane, targeting a certain physiological state is not more beneficial than simply reducing the dose.

### Physiological targets must be strain-specific

The significant effect of strain on network detectability was only revealed after physiological variables (in particular SpO_2_ and mean FD) were controlled for. C3HeB/FeJ have higher SpO_2_ by 6% and a lower FD by 1.87× 10^−3^mm on average than C57Bl/6 mice; these two characteristics bias them towards a good network detectability. However, if one were to compare a mouse from each strain with the same SpO_2_ and FD, the C3HeB/FeJ mouse would have significantly lower network detectability compared to the C57Bl/6 mouse (β=-0.15). Thus, SpO_2_ and FD are so-called suppressing mediators that mask the effect of strain. This finding underscores that it is risky to interpret physiological metrics from a new strain or model that one is not familiar with, as the same physiological state in two different strains can correspond to a different level of data quality.

Moreover, in all of the aforementioned regression pathways, there were significant interactions between anesthetic doses (isoflurane and dexmedetomidine) and demographic variables (strain and sex), indicating that network detectability and physiological variables were differentially affected by anesthetics depending on the strain and sex (Figures S4,S5).

### Network detectability declines over time

The significant relationship between time since isoflurane change and network detectability was also only revealed after controlling for mean FD. Later time windows contain less motion, which is generally associated with higher network detectability. However, comparing two time windows with equivalent motion reveals that network detectability decreases slightly over time (β=0.02), possibly indicating a deepening of anesthetic depth over time. If the linear trend continues, this effect would become noticeable by the end of a 20 minute scan.

### Dexmedetomidine has competing effects on network detectability

A reduced dexmedetomidine infusion of 0.05 mg/kg/h (compared to 0.1 mg/kg/h) was associated with improved SpO_2_ (β=-1.10) but also with increased FD (β=-0.431). The benefit of the increased SpO_2_ and the disadvantage of the increased FD canceled out such that dexmedetomidine had no significant net effect on network detectability. Nevertheless, further improvements to head restraint may control the increase in FD and justify the use of a reduced dexmedetomidine dose.

### Controlling isoflurane dose and SpO_2_ within a specific range yields a high success rate

We trained a decision tree algorithm to identify time windows that contain a recognizable network (i.e. whose network detectability is ≥ 0.45) from all other variables (anesthetic doses, session, physiology etc). The resulting decision tree corroborated the mediation analysis, with isoflurane dose again being the most important variables (Figure 3a), but it also determined the optimal range of values for physiological/motion variables in a data-driven approach and quantified the consequences of following a particular decision pathway. The decision tree was validated in a held-out test set of 112 time windows, exhibiting an accuracy of 79% (Figure 3b). Importantly, the rate of false positives was low, indicating that if the experimenter follows the decision paths that predict that a network will be present there is an 87% chance that they will indeed obtain a network.

**Figure 3:**
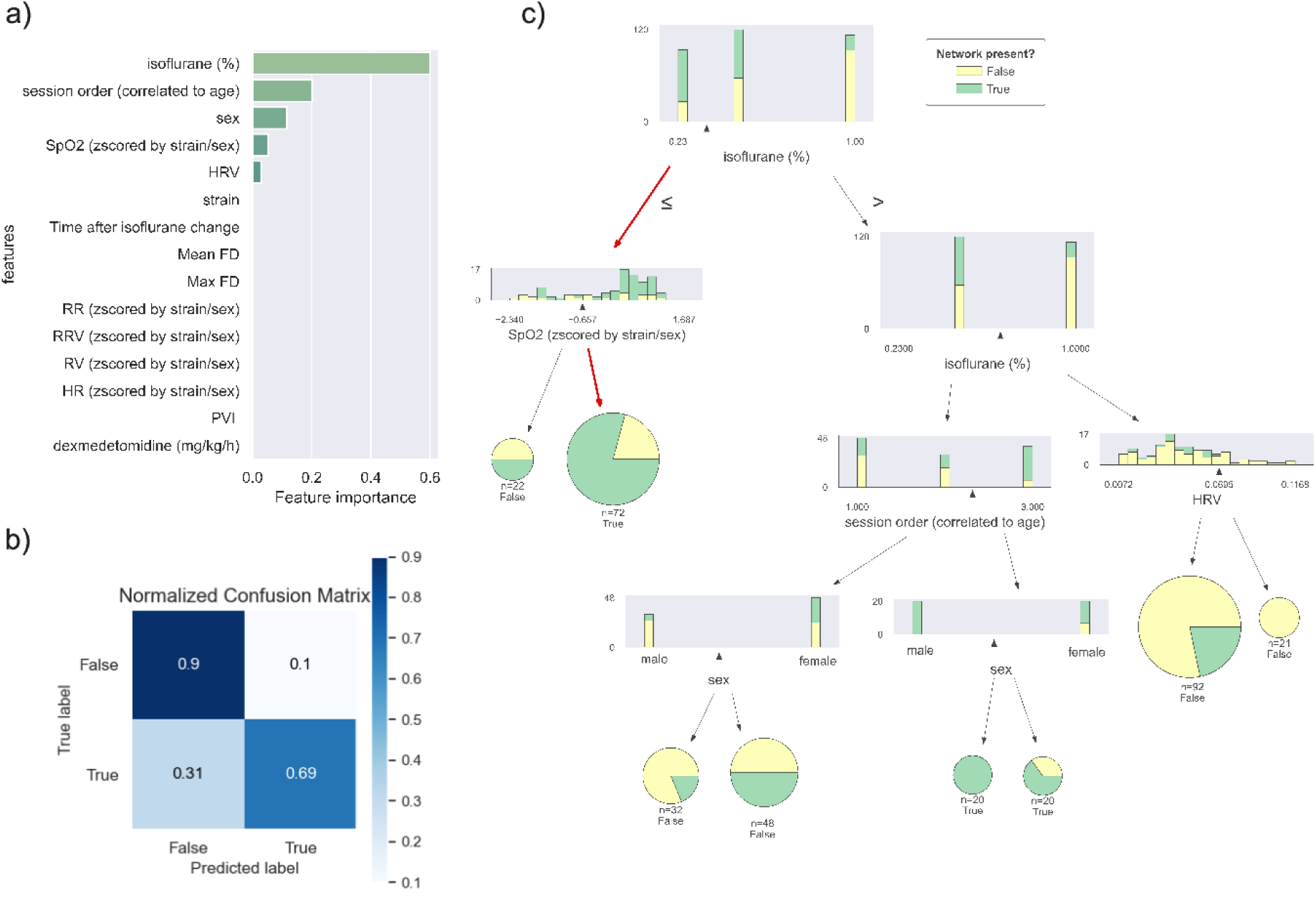
A decision tree identifies the acquisition choices that best separate time windows containing a network (network detectability > 0.45) from those that do not (network detectability <0.45). The decision tree is constructed layer-by-layer by splitting on the variable that will yield the purest leaves at each layer. a) The importance of each feature, computed as the mean decrease in Gini impurity index when splitting on that feature. All features that don’t appear in the final decision tree have an importance of 0. b) Confusion matrix obtained when the decision tree is applied to a held-out test set to classify time windows as containing a network or not, normalized by the number of samples (112 in the test set). c) The best path (i.e. the one that balances practicality and outcome success) is highlighted with red arrows: isoflurane should be 0.23% and aim for SpO_2_ values that are at least - 0.657 standard deviations above the strain-specific mean.

According to the decision tree, the ideal decision path that maximizes the likelihood that a time window contains a recognizable network is: use an isoflurane dose of 0.5% and scan mice two times before the main scanning session (or use mice that are 10-12 weeks old; the effect of age cannot be distinguished from the effect of session with our study design so it is not clear which one is driving the decision). Along this decision path, male mice will have a 100 % success rate and females will have a 65% success rate (Figure 3c).

Most study designs will not have the option of scanning mice twice beforehand (or of controlling age and sex). In that case, the next best decision path is to use an isoflurane dose of 0.23% and control SpO_2_ such that it is at least-0.7 standard deviations above the mean for a given strain, this corresponds to a minimal SpO_2_ value of 73% for C57Bl/6 mice and 86% for C3HeB/FeJ mice (Figure 3c, red arrows). This approach has a success rate of 81% on the level of 2-min time windows. On the level of 8-min time windows (i.e. a timescale that resembles more closely the duration of a normal scan), the scans acquired under these conditions had a success rate of 96% for the somatomotor network and 73% for the DMN-like network (Figure S6).

The decision tree also illustrates the consequences of following alternative paths. In particular, using an isoflurane dose of 1% will result in a high probability that the data contains no recognizable network, although odds are improved slightly at lower HRV values.

### Physiological states can be more useful than individual physiological metrics

Rather than individual physiological metrics, it may be more helpful to consider physiological states; namely co-varying patterns of respiration and plethysmography that recur across scans. To examine this possibility, we extracted 5 physiological states by performing K-Means clustering on the physiological metrics (after z-scoring within each strain/sex) (Figure 4a). We then trained a new decision tree, replacing the physiological metrics with the frequency of each physiological state within the 2-min time windows (Figure 4b).

**Figure 4.**
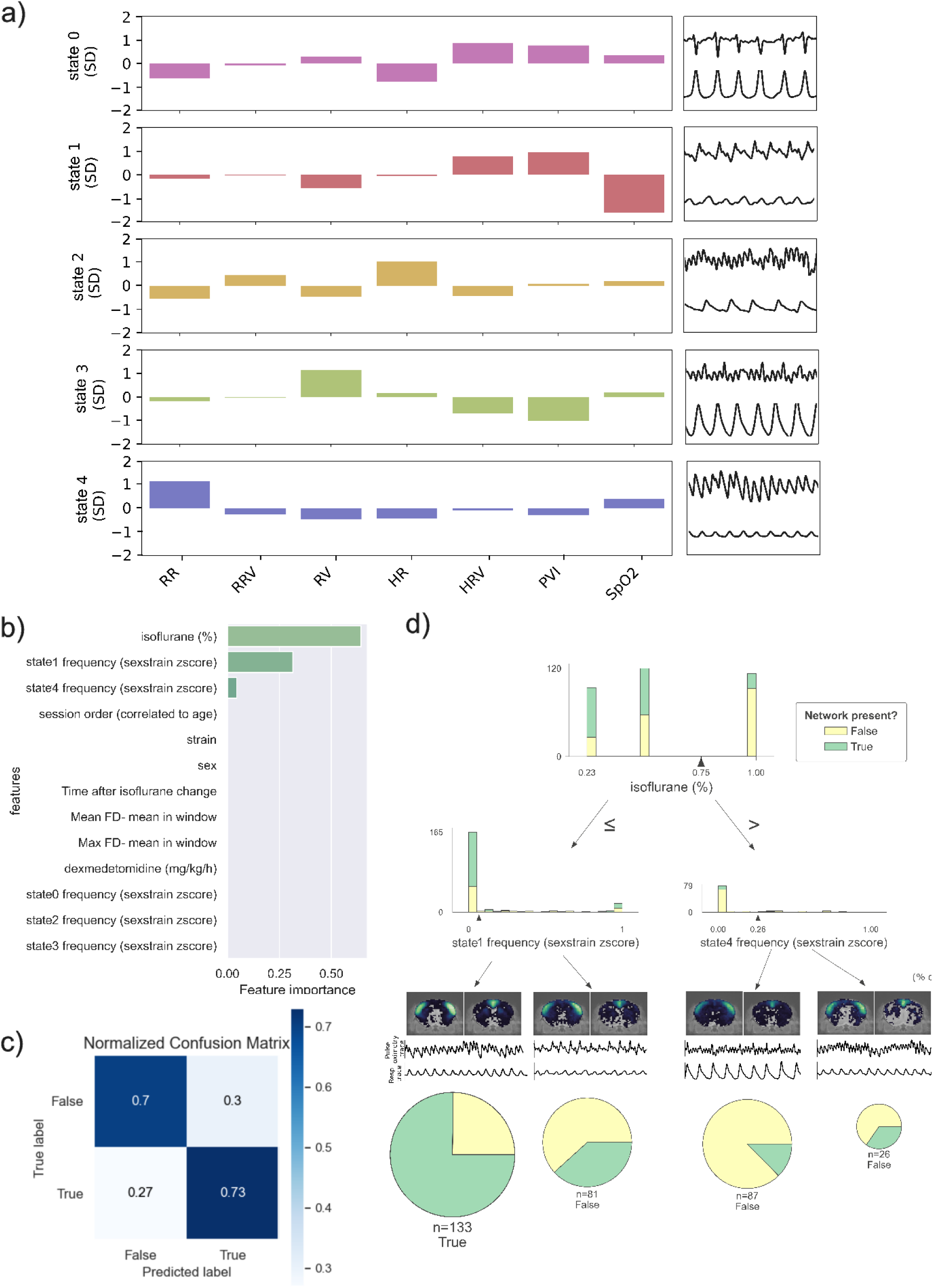
Considering the mouse’s global physiological state (i.e. the combined pattern of respiration and plethysmography) rather than individual metrics provides a more straightforward prediction of network detectability. a) The 5 physiological states obtained by K-Means clustering of the physiological metrics. The bar graph shows how each cluster centroid depends on each of the input physiological metrics - the y-axis is in standard deviations from the strain-specific mean since the physiological metrics were z-scored within strain before clustering. Next to the bar graph is a representative example of the plethysmography (upper) and respiration (lower) traces within each cluster. All examples were taken from C57Bl/6 mice. b) The importance of each feature, computed as the mean decrease in Gini impurity index when splitting on that feature. All features that do not appear in the final decision tree have an importance of 0. c) Confusion matrix obtained when the decision tree is applied to a held-out test set to classify time windows as containing a network or not, normalized by the number of samples. d) Decision tree obtained by training on physiological state variables instead of the individual physiological metrics. In this case, the best tree has only a depth of 2, and the ideal path consists of using isoflurane of 0.5% or below and avoiding the presence of physiological state #1. Frequency refers to the number of timepoints assigned to each physiological state within a given time window, thus a frequency of 0 means that that state never occurs within the time window and a frequency of 1 means that the entire time window consists only of that state. A representative example of the plethysmography and respiration traces in each of the final leaves of the tree is shown. The brain maps indicate the percentage of windows who have an active somatomotor (left) or DMN-like (right) network in a given voxel under each decision tree path.

The resulting tree had an out-of-sample accuracy of 71% (Figure 4c), slightly under that of the tree trained on the individual physiological metrics, however it was able to achieve this performance with a considerably shallower depth of 2 layers (Figure 4d). Based on this new tree, the ideal decision path when acquiring data would be to use an isoflurane dose of 0.5% or below and then avoid physiological state #1 consisting of low SpO_2_, irregular heart beats and shallow breathing.

Doing so would yield a success rate of 75% on network detectability. Thus, instead of controlling isoflurane precisely at 0.23% as in the previous section, there is more flexibility to use a higher isoflurane dose of 0.5% provided that abnormal state 1 is avoided. In both cases, maintaining good SpO_2_ levels is important for network detectability, but the physiological state results provide further context by indicating that low SpO_2_ is often accompanied by irregular heart beats and shallow breathing. Interestingly, HRV is high during state #1, which is an undesirable state, whereas the mediation analysis indicated that higher HRV was associated with improved network detectability. This underscores the importance of considering physiological values in the proper context: although high HRV can be indicative of alertness, it can also be indicative of poor oxygen saturation.

## Discussion

The need for accessible acquisition guidelines in the rodent rs-fMRI field is underscored by the recent publication of a standardized rat rs-fMRI “standardRat” protocol (Grandjean et al. 2023). However, there is no similar protocol for mice; past methodological publications for mouse fMRI have either focused solely on comparing anesthetics (Grandjean et al. 2014; Wu et al. 2017; Jonckers et al. 2014; Bukhari et al. 2017) or on combining fMRI with activation-based manipulations (optogenetic, pharmacological, task) (Adamczak et al. 2010; Ferrari et al. 2012; Petrinovic et al. 2016; Grimm, Wenderoth, and Zerbi 2022; Shim et al. 2022; Zeng et al. 2022). Therefore, as in the standardized rat protocol, the present study aimed to identify a set of accessible acquisition guidelines for mouse rs-fMRI that would aid researchers in attaining a baseline level of network detectability. Despite the overlapping goal, we employed a unique bottom-up approach that involved systematically varying anesthetic doses to probe different physiological/brain states (whereas standardRat was determined from a top-down comparison of many datasets). While both approaches have their own advantages, ours enables researchers to build a mechanistic understanding of the what (variables that have the strongest impacts), how (ideal parameter ranges) and why (whether variables have direct or indirect effects).

The success rate obtained with these recommendations is 93% for the somatomotor network and 73% for the DMN-like network (in 8-minute long scans), which is quite good for free-breathing, anesthetized mice. Note that the DMN is known to be harder to detect (Grandjean et al. 2023).

Previously, it was considered that mechanical ventilation was necessary for stabilizing physiology and obtaining high quality data - in fact, almost all of the aforementioned methodological publications for mouse fMRI use mechanical ventilation (Ferrari et al. 2012; Grandjean et al. 2014; Petrinovic et al. 2016; Wu et al. 2017; Bukhari et al. 2017; Grimm, Wenderoth, and Zerbi 2022; Shim et al. 2022). Given that ventilation is both challenging to implement (requiring intubation and sometimes paralysis of the mouse), requires specialized equipment, and can have negative consequences on the mouse’s health (De Vleeschauwer et al. 2011; Shim, Lee, and Kim 2020), our present demonstration of a successful free-breathing approach should be of great use to the field.

### Acquisition recommendations

#### RR and HR should not be used for tuning anesthetic dose

Mice undergoing anesthetic induction typically demonstrate dramatically deepening and slowing of their respiration due to the known effects of anesthesia on body physiology. As a result, multiple publications recommend using RR and HR as indicators of anesthetic depth (Constantinides, Mean, and Janssen 2011; Aline R. Steiner et al. 2020; Wallin et al. 2021; Navarro et al. 2021; Shim et al. 2022; Rivera et al. 2024) and even suggest adjusting anesthetic dose until a specific RR/HR is attained (Petrinovic et al. 2016; Reimann and Niendorf 2020). However, our results indicate that RR and HR are not predictive of network detectability above and beyond anesthetic dose itself. In other words, although both RR and network detectability decrease as isoflurane increases (in accordance with literature (Rivera et al. 2024)), RR doesn’t predict network detectability within a given dose - thus there can be no clear target RR to aim for. Similarly, HR increases with isoflurane (as in (Constantinides, Mean, and Janssen 2011; Grandjean et al. 2014) but doesn’t predict network detectability within a given dose. Essentially, RR/HR reflect anesthetic dose but there is no evidence that they reflect finer variables such as anesthetic depth or network detectability. Moreover, RR and HR are strongly dependent on strain and subject so attempting to interpret the values is risky. Therefore, although we did not measure anesthetic depth directly, we instead examined the end metric-of-interest, data quality (indexed via network detectability) and found both high and low quality data across a range of RR/HR values.

#### Choosing a sufficiently low isoflurane dose is the priority

Isoflurane dose had the single biggest impact on network detectability. Thus, rather than using physiology to adjust isoflurane during ongoing acquisition, researchers should prioritize piloting the lowest possible dose that is tolerated by their mouse model. In our mice, we found that 1% abolished networks, 0.5% yielded detectable networks and 0.23% increased detectability even further (note that these percentages refer to the values on our isoflurane vaporizer’s dial but the actual output may differ slightly depending on the calibration accuracy). The improvement in network detectability between 0.5% and 0.23% can be explained by increased HRV and FD; so in theory one could choose to adjust isoflurane within 0.23-0.5% using HRV and FD as markers (see next section). However, this is not strictly necessary; it is simpler to attempt the lowest possible isoflurane dose in all mice (although a certain percentage may wake up, 15% in our case). Within the literature, the standard isoflurane dose when using a combined isoflurane-(dex)medetomidine protocol is 0.5% (Grandjean et al. 2014, 2020) but a limited set of groups also chose to lower their dose to 0.2-0.3% (Pradier et al. 2021; Shim et al. 2022). Our finding leads to the question: would reducing isoflurane down to 0% (i.e. a dexmedetomidine-only protocol) yield even better results? Previous studies have shown that combining both anesthetics has multiple advantages including lower dose of either anesthetic, mitigated impacts on physiological parameters and improved network integrity. Overall, it is possible that isoflurane has a non-linear dose effect where the step from 0 to 0.23% differs from the step from 0.23% to 0.5% - indeed, HR under isoflurane follows a non-linear pattern where it decreases in comparison to awake but increases from lower to higher doses (Janssen et al. 2004; Constantinides, Mean, and Janssen 2011; Lairez et al. 2013). Thus, we recommend the combined isoflurane-medetomidine approach but with low levels of isoflurane.

#### SpO_2_ is an important indicator of an abnormal state and should be carefully monitored

Unusually low SpO_2_ (below 73% in C57Bl/6 and 86% in C3HeB/FeJ), while rare, is likely to yield poor network detectability. The exact underlying cause of such periods of low SpO_2_ are unclear - SpO_2_ is unrelated to isoflurane dose yet improves with decreasing dexmedetomidine as well as suddenly in the third session. The unclear origins make it hard to predict how to fix SpO_2_ in these cases, but possible options are: providing supplemental oxygen or reducing dexmedetomidine dose.

Note that according to literature, SpO_2_ <95% indicates that the mice are hypoxic, however all mice survived their scanning sessions, similarly to a previous publication reporting high survival rates despite apparent hypoxia with SpO_2_<70% (Blevins, Celeste, and Marx 2021).

#### HRV is a possible marker of anesthetic depth and may be useful to avoid awakening mice but should not be too heavily relied on

HRV predicts network detectability above and beyond anesthetic dose, implying that it may be sensitive to anesthetic depth - indeed HRV is frequently employed in the field of anesthesiology to evaluate loss of consciousness (Toweill et al. 2003; Mazzeo et al. 2011; W. N. P. Garcia 2021-2022). Thus, to lower the risk of mice awakening, researchers may monitor HRV and interpret higher values as a possible indication that the mouse is aroused and that it is not necessary to further lower anesthesia. However, this is tricky to do for multiple reasons: 1) there are no mouse physiological monitoring softwares to date that output real-time HRV, 2) elevated HRV also occurs during hypoxia (SpO_2_<70%) and must be interpreted in context of the SpO_2_ reading, 3) HRV depends strongly on subject and there is no specific threshold that can be relied upon as a general indicator across subjects. We are the first to measure HRV in the context of mouse fMRI and to propose it as a possible index of anesthetic depth.

#### Mild levels of motion need not be feared and should not be remedied by increasing isoflurane

FD also predicts network detectability above and beyond anesthetic dose: unsurprisingly, high FD (>0.01mm) predicts poor network detectability. However, mild motion below this threshold of 0.01mm is not correlated with network detectability hence researchers don’t need to focus on minimizing motion at all costs. If researchers experience problematic levels of motion, they should focus on improving head restraint rather than increasing isoflurane. The range of FD values displayed in the results does not include censored timepoints, thus the secondary consequences of motion spikes on timepoints after the movement itself were not considered. Note that our dataset contained relatively low levels of motion to begin with.

#### Physiological states may be useful indicators of network detectability

Our second decision tree indicated that a state characterized by simultaneously low SpO_2_ combined with elevated HRV and PVI was useful for predicting that networks would be abolished. Considering physiological state as a whole rather than individual metrics provides a more comprehensive understanding of the mouse’s condition and thus there is more flexibility to use either 0.23% or 0.5% isoflurane, providing that this state is avoided, without requiring a blanket solution of 0.23% isoflurane for all mice. However, in practice, considering physiological states is challenging because there is no real time classification algorithm for identifying the mouse’s state so it will have to be approximated by the experimenter, and secondly, it is unclear how to alter or fix this abnormal state. Moreover, we did not perform a mediation analysis with the physiological states so there may be additional context and interpretations that we lack.

### Study limitations and future directions

Despite our attempt to increase the generalizability of our results by including two strains, there are still many other strains that may respond differently. Thus, our findings are primarily a proof of concept that there are important differences between strains (particularly physiological ones) to encourage researchers to perform a pilot when using a strain/model that is new to them. Another demographic variable that we did not consider here is age although there is evidence that mammals become increasingly sensitive to anesthesia as they age (Eger 2001; Orliaguet et al. 2001; Drobac et al. 2004; Chemali et al. 2015). In summary, study designs with cross-strain comparisons or longitudinal trajectories, must take care to test that group differences or trajectories reflect actual differences in FC and not simply differences in anesthetic susceptibility or physiology.

Here, we focused on one anesthetic paradigm (isoflurane-dexmedetomidine combination) which tends to yield superior network detectability (Grandjean et al. 2023, 2020, 2014) and has become one of the most popular regimens; making it a great choice for newcomers to the field. However, isoflurane is known to have a narrow dose range between light and deep anesthetic depth, which makes it tricky to fine-tune anesthetic depth (Reimann and Niendorf 2020). Other promising anesthetics that have not been included in the major publications on anesthetic comparison (Williams et al. 2010; Grandjean et al. 2014; Jonckers et al. 2014; Paasonen et al. 2018) include halothane and etomidate. Thus, researchers that have difficulties implementing the isoflurane-medetomidine protocol (e.g. in novel mouse models) are encouraged to explore halothane and etomidate as options.

Session was another important predictor of network detectability but with our study design we were unable to definitively identify the explanation for this effect and turn it into an acquisition guideline. There was a sudden increase in network detectability during the third scanning session may be attributable to either: a) the development of chronic stress b) an effect of age, c) progressive hearing loss, d) habituation, e) cumulative effects of anesthesia exposure, f) an improvement in the experimenter’s ability to prepare the mouse. However, explanations a)-c) are unlikely since we observed normal weight gain, the sessions were only 3-10 days apart and mice were given ear plugs. Regardless, no single one of these explanations easily accounts for the fact that the session effect is not linear, with there being no difference between the first two sessions followed by a sudden improvement during the third. There may be multiple underlying causes contributing to the session effect and combining to produce the non-linear trajectory.

Finally, because the cause of inter-site variability in data quality is not yet fully understood (Grandjean et al. 2020), it remains to be seen to what extent our results reproduce in independent sites. We expect our main results to be robust (e.g. the dose-dependent impact of isoflurane on connectivity is well known, as reviewed in (A. R. Steiner, Rousseau-Blass, and Schroeter 2021)) however, the exact success rate that we obtain at each anesthetic dose may be different at other sites. In particular, differences in anesthetic susceptibility across sites may arise due to variations in mouse stress levels (L. Wang et al. 2019) and error margins or calibration differences between isoflurane vaporizers. For this reason, our guidelines are not written as a definitive protocol with specific parameters that may not generalize but are instead structured to help other researchers refine their own protocol.

### Open questions within the field

#### Which networks should be considered canonical and what is their ideal spatial representation?

Here, we defined ‘high’ quality data as having a specific spatial representation of the somatomotor and DMN-like networks. Within the literature, many other mouse resting-state networks have been described, but there is considerable variability in their granularity and the conditions under which they are detectable. The somatomotor is the most consistent network across all descriptions, thus we focus on it to provide a minimal baseline. It is possible that different acquisition recommendations might be more suited for detecting other networks. Moreover, even within the somatomotor and DMN-like networks, it is unclear whether a wider area of activation or a more localized one is “better”. That being said, our decision tree approach is independent of what the ideal spatial coverage is, as in both cases the network would be classified as detectable.

#### To what extent are fluctuations in network detectability across time windows attributable to biological variations in true network activity?

Such a question might be answerable with direct electrophysiological recordings of network activity in awake mice if one could be sure that there are no contributions of stress or motion. Indeed, all the variables that we could measure only explained 64% of the variance in network detectability, so it’s unclear what accounts for the remaining variance; it may be explained by the aforementioned biological changes in network activity, or variations in arousal or physiology (e.g. blood pressure, pCO_2_) that cannot be captured by simple physiological metrics.

*If reducing isoflurane dose is beneficial for network detectability, then is awake imaging even better?* There is little evidence in the literature that awake rs-fMRI data actually outperforms anesthetized data when it comes to network detectability (Yoshida et al. 2016), although awake mice do exhibit more complex network dynamics such as increased between-network communication (Gutierrez-Barragan et al. 2022). Thus, awake imaging may be more suitable for answering certain questions, but the benefits are not strong enough such that we would recommend it in general.

## Conclusions

In this work, we compared the relative importance of various acquisition-related variables (external ones such as anesthetic dose as well as continuous measurements of physiological parameters and motion) on network detectability in mouse rs-fMRI. We found that few continuous measurements can predict network detectability above and beyond the categorical external variables, and even those that can (i.e. HRV, mean FD and SpO_2_) are complicated to monitor in real-time and to subsequently interpret properly. Instead, we advise researchers using an isoflurane-(dex)medetomidine paradigm to prioritize reducing isoflurane dosage to the lowest level that doesn’t awaken their mice.

## Acknowledgements

Thank you to Alessandro Gozzi, members of the Gozzi lab, Tudor Ionescu and Yadunandan Edayadathodi for constructive criticisms and discussions.

## Notes

https://zenodo.org/uploads/12637274

https://github.com/CoBrALab/mousefmri_acq_publication/tree/main

https://github.com/CoBrALab/MousePhgyMetrics

https://github.com/CoBrALab/fMRI_phantom_analysis

### Competing Interest Statement

The authors have declared no competing interest.

