## Appendix for "Evidence-based guidelines for improving network detectability in rodent fMRI"

### Appendix: Guidelines for setting up an acquisition protocol for mouse fMRI

Refining acquisition is an iterative process that depends on the mouse model and equipment, thus we chose to provide 'guidelines' rather than a strict 'protocol'. We envision these guidelines being used in conjunction with the toolkits provided herein as well as our group's software for processing and quality control (RABIES) as follows: 1) the acquisition guidelines provide a starting point for pilot experiments, 2) RABIES is used to process, do basic confound correction, and quality control the pilot data, 3) if pilot data does not pass quality control, users can investigate possible sources of confounds, armed with the knowledge of their relative importance provided by the acquisition guidelines and/or the data quality assessment tools within RABIES (see details below). In this step, they can also make use of the toolkits for examining scanner stability or for quickly checking network detectability (in almost real-time) during the piloting session. It is recommended to avoid real-time checks of network detectability during the pilot and instead wait until this step, since attempting to tweak acquisition right from the first mouse may lead to overfitting and a biased intuition. 4) Finally, users can optimize their confound correction strategy to further improve data quality.

Notes:

- These guidelines are intended for newcomers to the field of mouse fMRI, thus we do not incorporate ventilation or awake imaging. We recommend starting with this free-breathing, minimally anesthetized approach then expanding to awake imaging if necessary.
- Even if the ultimate goal is not to study resting state networks per say, we regard the detection of activity from at minimum the somatomotor network as a necessary benchmark to ensure that one is not simply analyzing structured noise.

#### Step 1: Quality control the temporal stability of your MRI scanner

If any of the MRI scanner equipment contains temporal instabilities that manifest as changes in signal intensity over time, then your fMRI scans will be corrupted regardless of how well the mouse is prepared. Previous publications in the human literature have shown that faulty equipment can lead to major temporal instabilities that are not necessarily recognizable on structural scans (Friedman and Glover 2006). Do not assume that your MRI technician is performing regular quality control for temporal instabilities, this is not part of the standard procedures in the preclinical field.

I. Create an agar phantom that mimics the properties of the mouse brain,
   A. Combine 2% agarose powder (weight/volume) in 0.9% sodium chloride solution. It is also recommended to add 0.1M manganese chloride to better approximate the relaxation times of the mouse brain; a concentration of $2.0 \times 10^{-5}\%$ (volume/volume) manganese chloride is ideal for 7T scanners as it yields a T2* time of 43.5 ms, this concentration may need to be adjusted slightly for other field strengths. Optionally, 0.1% (weight/volume) of sodium azide can be added to prevent bacterial growth, this is recommended if the phantom will be used longitudinally.
   B. Slowly heat the mixture containing all ingredients in a microwave. Do so in 15-30 s intervals, stirring in between.
   C. Once the mixture turns into a clear liquid, fill a 3mL syringe (diameter of 9 mm, length of 6 cm) with the mixture.

D. Ideally, place the syringe in a vacuum to remove bubbles while the mixture is still liquid. Some number of bubbles will always persist.

E. Allow the mixture to cool. It will solidify into a gel.

II. Scan the agar phantom with the same fMRI sequence that you intend to scan mice with.

III. Process the scan with our toolkit ([https://github.com/CoBrALab/fMRI_phantom_analysis](https://github.com/CoBrALab/fMRI_phantom_analysis)) and follow the guidelines for interpretation. The principal component analysis (PCA) is particularly important to consider.

IV. If any metrics appear suspicious, alert your technician so that they can identify the cause. Depending on the magnitude, variance explained and spatial pattern of the temporal instability, it may still be possible to acquire good quality fMRI data despite its presence. This is a case by case assessment. For example, the IECO gradient amplifier clock used with certain Bruker scanners creates a sinusoidal instability, but since it has a very small amplitude relative to random noise, it is not a barrier to data quality.

V. Repeat steps II and III as part of a regular quality control procedure, at the very least prior to the start of each new fMRI project.

##### Step 2: Set up the required equipment for anesthesia and mouse monitoring.

I. Decide on the anesthetics to be used. The combination of isoflurane and medetomidine is commonly recommended as a balanced approach (Grandjean et al. 2014; Paasonen et al. 2018; Grandjean et al. 2020; A. R. Steiner, Rousseau-Blass, and Schroeter 2021). Note that medetomidine is a equal mixture dexmedetomidine (the active enantiomer) and levomedetomidine — hence, medetomidine and dexmedetomidine have the exact same mechanism of action except dexmedetomidine is twice as concentrated (Kuusela et al. 2001). Alternatively, halothane and etomidate are promising anesthetics that have not been included in formal published anesthesia comparisons. Halothane has been overlooked largely due to its hepatotoxicity, but there is little evidence that the hepatotoxic effect is significant in mice at the moderate doses used for fMRI (even in humans the initial exposure is not clinically significant (Elliott and Strunin 1993)). Certain groups endorse the use of halothane for rodent rs-fMRI due to its ability to produce clean networks (Sforazzini et al. 2014; Liska et al. 2015) and its wide dose range (Reimann and Niendorf 2020). Unfortunately, as of January 2025 there is a global shortage of halothane. Etomidate on the other hand, is unique because it is an injectable anesthetic that has minimal impacts on hemodynamics, produces consistent responses across strains and is suitable for longitudinal experiments (Petrinovic et al. 2016) yet despite these benefits has rarely been used in studies.

II. Create a setup for delivering oxygen-enriched air to the mice. Regular medical air is insufficient since mice become hypoxic under anesthesia and thus require supplemental oxygen (Lumb 2019). Pure 100% oxygen, as is sometimes used in structural imaging protocols, is also not ideal as it may induce hypoventilation and alter the baseline cerebral blood flow (CBF) and BOLD signals (Sicard and Duong 2005). To best reproduce a normal homeostatic state, it is common to use an air-to-oxygen ratio between 80:20 and 60:40 (Baltes et al. 2011; Fukuda et al. 2013; Grandjean et al. 2014; Zerbi et al. 2015; Brynildsen et al. 2017).

III. Create a setup for the delivery of isoflurane (or other volatile anesthetics).

A. Confirm that the isoflurane machine has been calibrated within the last year. Beware that isoflurane machines are usually calibrated to be within ±20% of the marked values (0.5%, 1%, 2%, 3%, 4%, 5%) as they are intended for use during surgeries. A 20% error margin is already large, but they may be even less accurate at low values below 0.5%. Moreover, isoflurane output can drift over time and a minor drift (say from 0.3% to 0.5%) may not be noticed during calibration but will nevertheless impact the mice. It is recommended to either buy an isoflurane meter or request the isoflurane machine calibration technician to check the output within the low range as well.

B. Confirm that the isoflurane is properly exiting the nose cone and that the waste gases are being scavenged. This is particularly important if you have a custom built nose cone. If the expired gases build up in the nose cone, the mouse will re-breath them and may become hypercapnic. Buildup can also result in the effective isoflurane dose changing over time.

C. Confirm that the mouse model that you are using fits snugly into the nose cone and bite bar. Certain mouse strains can have abnormal tooth lengths or nose shapes, hence it is recommended to check this during a pilot when starting a study with a new mouse strain.

IV. Decide on the delivery route for injectable anesthetics. In the case of (dex)-medetomidine, multiple routes have been proposed, including subcutaneous (Adamczak et al. 2010; Jonckers et al. 2011; Petrinovic et al. 2016; Pradier et al. 2021), intraperitoneal (Nasrallah et al. 2014) or intravenous (Grandjean et al. 2014; Gutierrez-Barragan et al. 2022). The intravenous approach has been shown to yield more stable FC patterns during long imaging sessions compared to subcutaneous (Sirmpilatze, Baudewig, and Boretius 2019). Regardless, all of these approaches recommend an initial bolus followed by a continuous infusion (via an infusion pump).

V. Check the shelf life of all your anesthetics. In particular, medetomidine hydrochloride solutions (e.g. Domitor) have a short shelf life of 3 months after opening and their efficacy at sedation may decay if it is used past that.

VI. Develop a setup for controlling and monitoring mouse physiology.

A. While anesthetized, thermoregulation is impaired, hence it is necessary to maintain the mouse within a normal temperature range (~36.5℃) artificially. Typically, this is achieved by monitoring internal temperature with a rectal thermometer and heating the mouse accordingly via a warm water bed or air heater that connects to the thermometer with a feedback loop. A feedback loop for tight control is crucial since even small changes in body temperature (<1℃) can cause changes in BOLD signal intensity (Vanhoutte, Verhoye, and Van der Linden 2006). Some sort of heating even during induction would be ideal, as body temperature drops very quickly - it has been reported that the BOLD response can be impacted even after body temperature returns to normal following mild hypothermia (Shim et al. 2022).

B. Respiration is monitored with a respiration pillow while heart rate and oxygen saturation are monitored with an ankle pulse oximeter. We recommend the ankle pulse oximeter (https://i4sa.com/product/pulse-oximeter-fo-sensor-ankle-tail-form-mouse/) over a clip-on pulse oximeter as it does not restrict blood flow. We find that the plethysmography trace frequently looks noisy during the mouse setup but it stabilizes over time after the mouse is placed in the scanner. Of these metrics, the most important to monitor is $SpO_2$.

VII. Develop a setup for head fixation. Minimal head motion is important for preventing motion-related changes in signal intensity, distortions and signal dropout. Note that although EPI frames can be realigned during processing, this does not correct for the secondary consequences of motion such as the spin-history effect (Murphy, Birn, and Bandettini 2013). Therefore, motion during acquisition needs to be below a certain threshold, although too little motion can be a sign of over anesthesia and is actually associated with poorer network detectability. As such, motion should be prevented via head fixation and not by over-anesthetizing. Various options are possible, including ear bars, custom nose cones and head plate implants. We have found that the ear bar setup that is built into the Bruker bed is not sufficiently adjustable for each mouse and does not provide the desired level of head fixation. We prefer to use padding between the mouse's head and the CryoProbe™ surface coil such that there is little space for the head to move - the appropriateness of this approach will depend on the specific coil and mouse configuration.

##### Step 3: Acquire rs-fMRI data

I. Prepare the solutions in advance if using injectable anesthetics such as (dex)medetomidine. We use a dexmedetomidine bolus of 0.05 mg/kg (intraperitoneal) and a dexmedetomidine infusion of 0.025 mg/kg/h (intraperitoneal). If using (dex)medetomidine, it is recommended to also inject atipamezole to reverse the effects of (dex)medetomidine after the scan - atipamezole is 10 times less potent than dexmedetomidine and 5 times less potent than medetomidine. Note that standard injection procedures recommend injecting no more the 0.01 mL per gram of mouse weight, so choose a dilution ratio accordingly. We use a 1:60 dilution of 0.5mg/mL dexmedetomidine (final concentration of 0.0083 mg/mL) and 1:20 dilution of 5mg/mL atipamezole (final concentration of 0.25 mg/mL).

II. Prepare the equipment: nose cone, respiration pillow, pulse oximeter, heater, temperature probe, syringe and pump for continuous infusion*, isoflurane and oxygen-enriched air delivery, ear plugs, head padding.

III. Induce anesthesia.
   A. Weigh the mouse
   B. Calculate injection volumes*
   C. Induce anesthesia with 3.5% isoflurane and wait until the mouse has noticeably slower and deeper breathing (~5 minutes). Ideally, keep the mouse warm throughout the induction.
   D. Inject (dex)medetomidine bolus*. Can be done intraperitoneally (Nasrallah, Tay, and Chuang 2014), subcutaneously (Adamczak et al. 2010; Jonckers et al. 2011; Petrinovic et al. 2016; Pradier et al. 2021) or intravenously (Grandjean et al. 2014; Gutierrez-Barragan et al. 2022). It may be desirable to adjust the timing of future steps based on the injection method used. In this study, we used intraperitoneal injections for convenience. We provide example calculations for injection volume (for ip injection of dexmedetomidine using a 1:60 dilution of dexmedetomidine).

$$dex\ bolus\ volume\ (mL) = \frac{0.05\ (mg/kg) * mouse\ weight\ (kg)}{0.0083\ (mg/mL)}$$

$$dex\ infusion\ rate\ (mL/h) = \frac{0.025\ (mg/kg/h) * mouse\ weight\ (kg)}{0.0083\ (mg/mL)}$$

   E. Apply ophthalmic ointment to prevent eye dryness.
   F. Replace the mouse in the induction chamber at 2% isoflurane until they are re-anesthetized.

IV. Set up the mouse on the scanner bed.
   A. Rapidly insert the needle for continuous infusion intraperitoneally (Figure A1.2a).
   B. Place the mouse on the scanner bed on top of the respiration pillow, and secure them in the nose cone (Figure A1.2b,c). The nose cone should be delivering 1.5% isoflurane in a 20:80 oxygen to air mixture (up to 40:60 is acceptable).

C. Check that the mouse is properly secured in the nose cone by tugging gently backwards near the ears. The nose cone should not appear painfully tight and breathing pattern should not change before and after the nose cone is tightened.

D. Place a disposable cover on the rectal temperature probe and cover it with vaseline. Insert the probe to the same depth each time. Start heating immediately.

E. Take small pieces of the ear plug putty (e.g. Mack's silicone ear plugs), roll into ovals and insert into the mouse's ears. The ear plug should be small enough so that it doesn't require pushing into the ears. (Figure A1.2d)

F. Use tweezers to place the mouse's ankle inside the pulse oximeter. Make sure all the toes are inside. Then tape the pulse oximeter tubes (don't tape over the opening where the light is). The final oximeter position should be on the lower part of the ankle, right near the foot. (Figure A1.2e)

G. Double check that the head is still secure inside the nose cone and that it is not tilted within the cone. Add the padding on top of the head (if using a porous material, it should be replaced for each mouse). (Figure A1.2f)

H. Confirm that the respiration rate and heart rate are being properly recorded. We have noticed that the pulse oximetry signal looks noisy initially but often stabilises a few minutes after the mouse is placed in the scanner.

I. The dexmedetomidine infusion can be started at 10-15 minutes after the original bolus. This suggestion is an estimate - if the mice are consistently waking up at a certain time following the bolus injection, it may be due to depletion of the bolus effect, in which case try starting the infusion sooner (this should be determined during piloting).

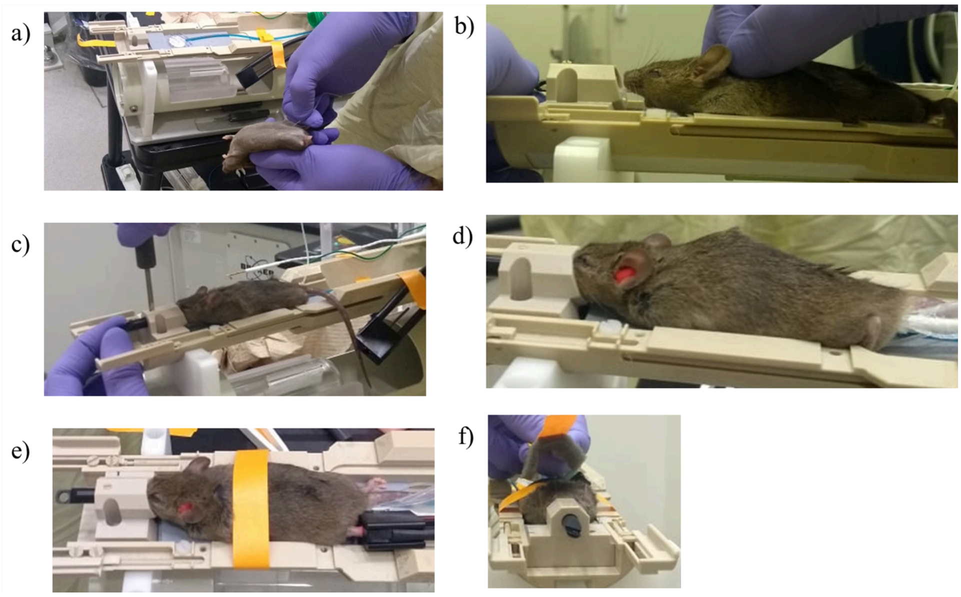


***Figure A1.2:*** *Steps for setting up the mouse on the scanner bed. a) Injecting the infusion needle. b) Placing the mouse into the nose cone. c) Tightening the nose cone. d) Ear plugs and temperature probe have been inserted. e) Ankle pulse oximeter has been placed. f) Checking head tilt and placing padding.*

V. Scanning.

A. Perform B0 mapping and shimming over the region of interest. This is important for reducing the distortions and signal drop out that EPI scans are particularly prone to.

B. If acquiring high-resolution anatomical scans that will be further analyzed (i.e. not just to be used as a reference for the functional scans), do this early while isoflurane is still high (at least 1%). Acquiring anatomical scans at lower doses may result in motion artifacts. The longer the anatomical scan is, the more prone it is to motion artifacts, since even a single movement will impact the whole scan and cannot be censored afterwards (unlike in fMRI).
C. Gradually reduce isoflurane dose until the desired final concentration (0.5% at most) is reached. Physiological parameters (like $SpO_2$) will gradually improve during the reduction. The exact turn down procedure is not important, so long as there is a minimum of 5 minutes between turn downs for the mice to partially stabilize, and the turn downs are decreases of 0.2-0.5% each time. An example is provided (figure A1.1).
D. Wait at least 10 minutes at the final dose for the mouse to stabilize. This should be at least 45 minutes after the bolus injection (Pradier et al. 2021).
    - If the $SpO_2$ is still below 73% (for C57Bl/6 mice, or 85% for C3HeB/FeJ mice), increase the ratio of oxygen to air delivered or try decreasing isoflurane further.
      **Note that mice (and humans) experience neural inertia when emerging from anesthesia, this means that they can get trapped in an unconscious state. Thus, it may be necessary to substantially lower isoflurane until they escape this state of deeper anesthetic depth (often visible as a sudden change in physiology) then increase it back to the normal dose. When increased back up, they will not revert to the state of deeper depth since the transitions into and out-of anesthesia exhibit hysteresis. The goal is not for the mouse to wake up but simply escape a state of over-anesthesia into lighter anesthesia. This process of stepping down then back up for some mice that appear to be over-anesthetized is a convenient way to acquire all the mice at the same final dose.
    - There should be indicators of minor motion at this dose that are noticeable as sudden voltage changes on the respiration trace or in the camera.
    - Wait again for the mouse to stabilize after any changes.
E. Acquire the EPI sequence.
F. If desired, check the quality of the acquisition with a quick ICA before removing the mouse using this script that is designed for quick on-the-spot checks: (https://github.com/Gab-D-G/quick_QC). If you are doing piloting, it is suitable to use this feedback to try tuning dosage in order to establish a final protocol, just bear in mind that network detectability decreases slowly over time so long piloting sessions in a single animal may yield biased conclusions. If you are acquiring for the main experiment, tuning dosage on a per mouse basis may be controversial.
G. Inject atipamezole after removing the mouse. They should appear fully recovered and moving within 10 minutes of the atipamezole injection.

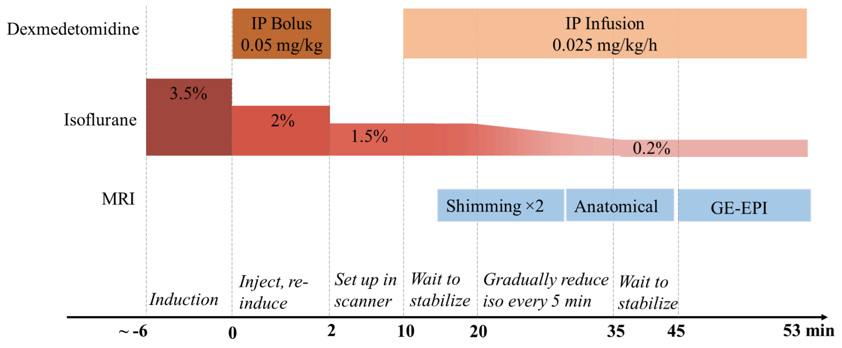


***Figure A1.1:*** *Diagram of anesthesia timing.*

##### Step 4: Confirm the quality of your dataset using the RABIES quality control tools

The RABIES software includes a suite of automatically-generated data quality reports that allow for estimating both network detectability and potential sources of confounds in the analysis, and are generally helpful in characterizing potential issues. The details and guidelines for conducting quality control using those reports are documented online (https://rabies.readthedocs.io/en/latest/analysis_QC.html) and in the original software publication (Desrosiers-Grégoire et al. 2024). These reports can assist with making informed changes to the acquisition protocol as follows: 1) very low network detectability values may indicate that the anesthetic dose is too high, 2) frequent spikes in FD that induce changes in global signal may indicate that head restraint approach needs to be improved (see figure S4 from (Desrosiers-Grégoire et al. 2024)), 3) slow fluctuations in global signal that appear as vascular artifacts may indicate problems with physiology (see figure S5 from (Desrosiers-Grégoire et al. 2024)) - in this case, if anesthetic is already low then carefully reread this appendix to be sure other aspects are not neglected. Note that these aforementioned points are only problematic if they appear in a significant number of mice, individual mice may have artifacts, you should not adjust your whole protocol in an attempt to achieve a 100% success rate.

RABIES is also helpful after the protocol has been finalized to determine the number of mice needed in the final experiment, depending on the percentage of scans that exhibit desired network detectability with the current protocol. In particular, RABIES generates distribution plots (https://rabies.readthedocs.io/en/latest/nested_docs/distribution_plot.html) that indicate the presence of networks and correlation with confounds at the scan level, which are useful for identifying outliers in the dataset and scans that do not meet the minimum quality criteria. Figure S6 is similar to these plots and can serve as a reference.
