## Supplementary for "Evidence-based guidelines for improving network detectability in rodent fMRI"

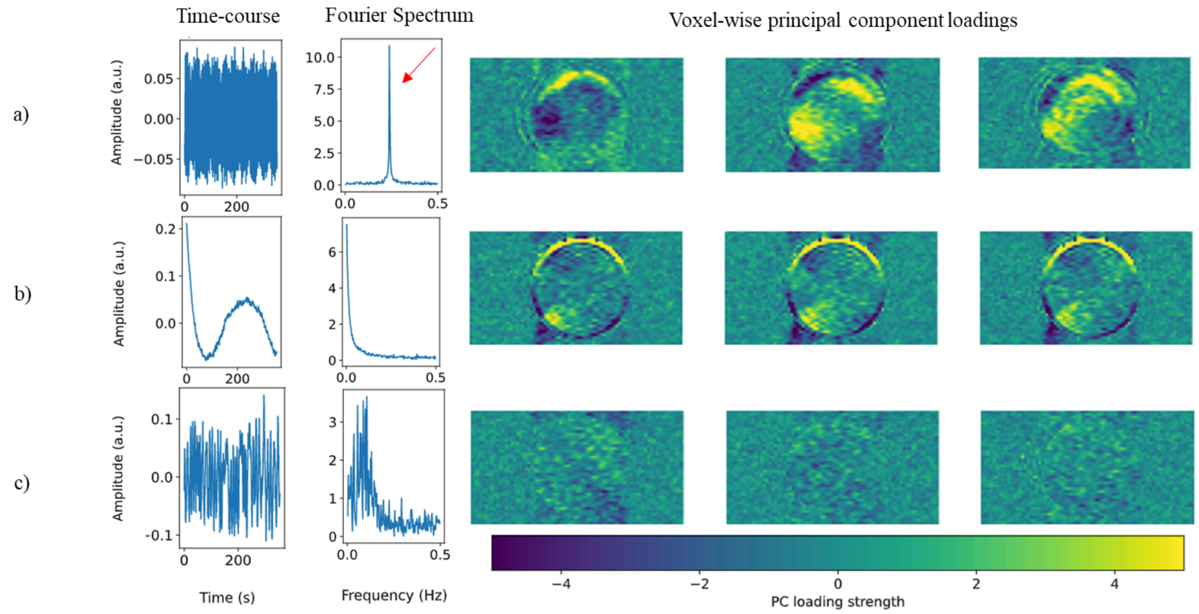


*Figure S1. Three different principal components from the same fMRI run, outputted by the temporal stability quality assessment (QA) report from our custom toolkit [https://github.com/CoBrALab/fMRI_phantom_analysis](https://github.com/CoBrALab/fMRI_phantom_analysis). a) component depicting periodic instability (explains 4.1% of variance), b) component depicting the slowly-varying cubic instability (explains 4.7% of variance), c) component depicting random noise (explains 1.1% of the variance) — the remaining components all resemble this one.*

a) Means

| | SpO2 | PVI | HR | HRV | RV | RR | RRV |
|---|---|---|---|---|---|---|---|
| **C57Bl/6 males** | 82.101 | 83.69 | 290.054 | 0.052 | 8.675 | 102.592 | 0.033 |
| **C57Bl/6 females** | 81.697 | 73.816 | 306.561 | 0.036 | 2.751 | 126.5 | 0.02 |
| **C3HeB/FeJ males** | 91.227 | 69.831 | 208.943 | 0.057 | 8.545 | 84.714 | 0.037 |
| **C3HeB/FeJ females** | 87.014 | 83.054 | 236.676 | 0.055 | 7.507 | 84.38 | 0.05 |

b) Standard Deviations

| | SpO2 | PVI | HR | HRV | RV | RR | RRV |
|---|---|---|---|---|---|---|---|
| **C57Bl/6 males** | 14.651 | 12.491 | 39.07 | 0.027 | 4.323 | 16.114 | 0.032 |
| **C57Bl/6 females** | 9.668 | 17.63 | 41.372 | 0.036 | 1.454 | 14.591 | 0.021 |
| **C3HeB/FeJ males** | 2.875 | 14.894 | 18.926 | 0.026 | 3.091 | 8.338 | 0.027 |
| **C3HeB/FeJ females** | 6.179 | 10.542 | 27.935 | 0.033 | 2.603 | 11.521 | 0.055 |

*Figure S2. The mean (a) and standard deviation (b) of each physiological metric within the strain and sex subgroups. The physiology data was z-scored within each strain and sex to remove the differences across strain/sex in order to construct an accurate decision tree and for the purpose of plotting the mediation analysis results.*

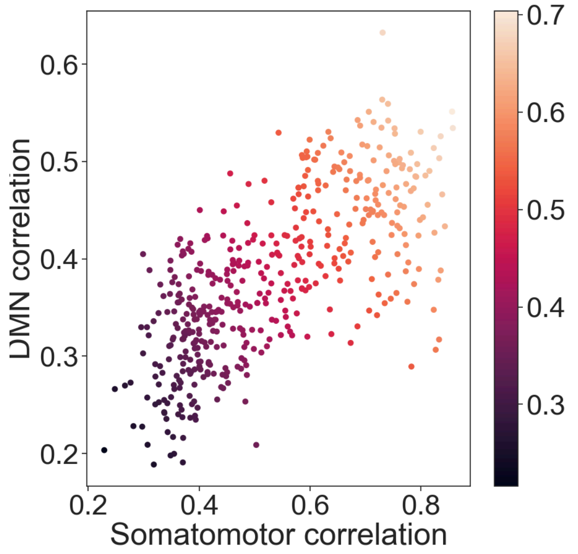


*Figure S3. Relationship between the detectability of the somatomotor network and the detectability of the DMN-like network across 2-minute time windows. The color bar indicates the average detectability of the two networks - this is the final metric that is referred to as 'network detectability' throughout the study. The correlation values on the x and y axis refer to the spatial correlation between the DR outputs and the original, group-level network map. Overall, time windows that contain one network often contain the other as well, there is just a small subset of windows with high somatomotor detectability where the DMN is hard to detect.*

| **Mouse (sex, strain)** | **Session number (dexmedetomidine infusion)** | **Timepoints excluded** | **Reason** |
|---|---|---|---|
| 006 (f, C57Bl/6) | 3 (0.05 mg/kg.h) | Full scan | Mouse died between the second and third scanning session. Reason unknown. |
| 009 (m, C3HeB/FeJ) | 2 (0.05 mg/kg/h) | Full scan | Mouse moved out of FOV during scanning and it was not noticed until later. |
| 016 (f, C3HeB/FeJ) | 3 (0.025 mg/kg/h) | 1380-1440 | Too much motion censoring. |
| 001 (m, C57Bl/6) | 2 (0.05 mg/kg/h) | Full scan | Pulse oximetry data didn't pass QC; beats could not be reliably identified. |
| 003 (m, C57Bl/6) | 1 (0.025 mg/kg/h) | 475-1440 | Pulse oximeter fell off (or was kicked off). |
| 004 (m, C57Bl/6) | 2 (0.1 mg/kg/h) | Full scan | Pulse oximetry data didn't pass QC; beats could not be reliably identified. |
| 004 (m, C57Bl/6) | 1 (0.05 mg/kg/h) | 1120-1440 | Pulse oximetry data didn't pass QC; beats could not be reliably identified. |
| 005 (f, C57Bl/6) | 2 (0.05 mg/kg/h) | Full scan | Pulse oximetry data didn't pass QC; beats could not be reliably identified. |

| | | | |
|---|---|---|---|
| 013<br>(f, C3HeB/FeJ) | 2<br>(0.05 mg/kg/h) | 1380-1440 | Pulse oximetry data didn't pass QC; beats could not be reliably identified. |
| 007<br>(f, C57Bl/6) | 3<br>(0.1 mg/kg/h) | Full scan | Respiration data didn't pass QC; breaths could not be reliably identified. |
| 008<br>(f, C57Bl/6) | 2<br>(0.1 mg/kg/h) | 0-450 | Respiration data didn't pass QC; breaths could not be reliably identified. |
| 010<br>(m, C3HeB/FeJ) | 2<br>(0.025 mg/kg/h) | Full scan | Physiological rates and SpO2 trace not available. |

*Table S1. Data excluded from the final analysis due to quality control or attrition.*

Table S1: Dependence of mediators (physiology, motion) on independent variables (demographics, anesthesia, session, time)

Path A of mediation analysis

| Mediator Variable | Independent Variable | Effect estimate (standardized) | Confidence Interval (lower) | Confidence Interval (upper) | Significant? |
|---|---|---|---|---|---|
| RR | strain (C3HeB/FeJ) | -1.573 | -2.039 | -1.107 | * |
| RR | sex (f) | 0.423 | -0.014 | 0.884 | |
| RR | isoflurane (0.5% vs 0.23%) | -0.237 | -0.415 | -0.060 | * |
| RR | isoflurane (1% vs 0.5%) | -0.594 | -0.765 | -0.421 | * |
| RR | dexmedetomidine (0.05 vs 0.025) | -0.866 | -1.072 | -0.658 | * |
| RR | dexmedetomidine (0.1 vs 0.05) | 0.892 | 0.681 | 1.106 | * |
| RR | session (2 vs 1) | -0.087 | -0.199 | 0.031 | |
| RR | session (3 vs 2) | -0.071 | -0.200 | 0.064 | |
| RR | Time after isoflurane change | -0.098 | -0.139 | -0.058 | * |
| RRV | strain (C3HeB/FeJ) | 0.705 | 0.242 | 1.149 | * |
| RRV | sex (f) | 0.290 | -0.177 | 0.740 | |
| RRV | isoflurane (0.5% vs 0.23%) | -0.041 | -0.387 | 0.286 | |
| RRV | isoflurane (1% vs 0.5%) | 0.181 | -0.142 | 0.504 | |
| RRV | dexmedetomidine (0.05 vs 0.025) | 0.636 | 0.241 | 1.039 | * |
| RRV | dexmedetomidine (0.1 vs 0.05) | -0.945 | -1.327 | -0.549 | * |
| RRV | session (2 vs 1) | -0.259 | -0.481 | -0.038 | * |
| RRV | session (3 vs 2) | 0.643 | 0.395 | 0.887 | * |
| RRV | Time after isoflurane change | 0.052 | -0.023 | 0.129 | |
| RV | strain (C3HeB/FeJ) | 0.644 | 0.073 | 1.243 | * |
| RV | sex (f) | -0.972 | -1.559 | -0.404 | * |
| RV | isoflurane (0.5% vs 0.23%) | -0.256 | -0.441 | -0.064 | * |
| RV | isoflurane (1% vs 0.5%) | 0.053 | -0.126 | 0.225 | |
| RV | dexmedetomidine (0.05 vs 0.025) | 1.299 | 1.074 | 1.530 | * |
| RV | dexmedetomidine (0.1 vs 0.05) | -1.237 | -1.456 | -1.009 | * |
| RV | session (2 vs 1) | 0.835 | 0.709 | 0.961 | * |
| RV | session (3 vs 2) | -0.210 | -0.343 | -0.075 | * |
| RV | Time after isoflurane change | -0.028 | -0.068 | 0.013 | |
| HR | strain (C3HeB/FeJ) | -1.399 | -1.965 | -0.827 | * |
| HR | sex (f) | 0.483 | -0.094 | 1.056 | |
| HR | isoflurane (0.5% vs 0.23%) | 0.379 | 0.186 | 0.574 | * |
| HR | isoflurane (1% vs 0.5%) | 0.519 | 0.330 | 0.706 | * |
| HR | dexmedetomidine (0.05 vs 0.025) | 0.210 | -0.033 | 0.445 | |
| HR | dexmedetomidine (0.1 vs 0.05) | -0.652 | -0.889 | -0.417 | * |
| HR | session (2 vs 1) | 0.109 | -0.019 | 0.240 | |
| HR | session (3 vs 2) | -0.265 | -0.408 | -0.127 | * |
| HR | Time after isoflurane change | 0.021 | -0.023 | 0.064 | |

The effect of independent variables on mediator variables, modeled as a linear mixed effects regression using Bayesian statistics. All mediator variables are continuous and standardized to a mean of 0 and standard deviation of 1. Thus the effect estimate indicates the mean difference in the mediator variable (eg RR) when comparing the two contrasts (e.g. two strains), expressed in units of standard deviation. The 95% confidence intervals indicate the lowest and highest esimates for the effect size. If the confidence interval crosses 0, then the effect is not significant.

Table S1: Dependence of mediators (physiology, motion) on independent variables (demographics, anesthesia, session, time)

Path A of mediation analysis

| Mediator Variable | Independent Variable | Effect estimate (standardized) | Confidence Interval (lower) | Confidence Interval (upper) | Significant? |
|---|---|---|---|---|---|
| PVI | strain (C3HeB/FeJ) | 0.071 | -0.721 | 0.852 | |
| PVI | sex (f) | 0.414 | -0.364 | 1.181 | |
| PVI | isoflurane (0.5% vs 0.23%) | -0.133 | -0.392 | 0.136 | |
| PVI | isoflurane (1% vs 0.5%) | 0.082 | -0.165 | 0.332 | |
| PVI | dexmedetomidine (0.05 vs 0.025) | 1.370 | 1.065 | 1.692 | * |
| PVI | dexmedetomidine (0.1 vs 0.05) | -0.735 | -1.049 | -0.431 | * |
| PVI | session (2 vs 1) | -0.269 | -0.435 | -0.099 | * |
| PVI | session (3 vs 2) | 0.498 | 0.309 | 0.688 | * |
| PVI | Time after isoflurane change | 0.014 | -0.045 | 0.071 | |
| HRV | strain (C3HeB/FeJ) | 0.559 | -0.329 | 1.447 | |
| HRV | sex (f) | -0.215 | -1.109 | 0.680 | |
| HRV | isoflurane (0.5% vs 0.23%) | -0.339 | -0.563 | -0.112 | * |
| HRV | isoflurane (1% vs 0.5%) | -0.303 | -0.521 | -0.089 | * |
| HRV | dexmedetomidine (0.05 vs 0.025) | 0.506 | 0.234 | 0.774 | * |
| HRV | dexmedetomidine (0.1 vs 0.05) | 0.187 | -0.080 | 0.456 | |
| HRV | session (2 vs 1) | -0.137 | -0.286 | 0.011 | |
| HRV | session (3 vs 2) | 0.316 | 0.153 | 0.478 | * |
| HRV | Time after isoflurane change | -0.010 | -0.058 | 0.040 | |
| SpO2 | strain (C3HeB/FeJ) | 0.618 | 0.141 | 1.100 | * |
| SpO2 | sex (f) | -0.195 | -0.697 | 0.303 | |
| SpO2 | isoflurane (0.5% vs 0.23%) | -0.072 | -0.350 | 0.207 | |
| SpO2 | isoflurane (1% vs 0.5%) | 0.000 | -0.260 | 0.261 | |
| SpO2 | dexmedetomidine (0.05 vs 0.025) | -0.187 | -0.513 | 0.128 | |
| SpO2 | dexmedetomidine (0.1 vs 0.05) | -1.096 | -1.413 | -0.772 | * |
| SpO2 | session (2 vs 1) | -0.043 | -0.223 | 0.142 | |
| SpO2 | session (3 vs 2) | 0.894 | 0.691 | 1.095 | * |
| SpO2 | Time after isoflurane change | 0.011 | -0.050 | 0.073 | |
| MeanFD | strain (C3HeB/FeJ) | -0.762 | -1.270 | -0.266 | * |
| MeanFD | sex (f) | -0.187 | -0.729 | 0.346 | |
| MeanFD | isoflurane (0.5% vs 0.23%) | 0.053 | -0.257 | 0.389 | |
| MeanFD | isoflurane (1% vs 0.5%) | -0.359 | -0.662 | -0.047 | * |
| MeanFD | dexmedetomidine (0.05 vs 0.025) | 0.028 | -0.345 | 0.405 | |
| MeanFD | dexmedetomidine (0.1 vs 0.05) | -0.431 | -0.788 | -0.050 | * |
| MeanFD | session (2 vs 1) | -0.388 | -0.590 | -0.181 | * |
| MeanFD | session (3 vs 2) | -0.422 | -0.646 | -0.196 | * |
| MeanFD | Time after isoflurane change | -0.080 | -0.150 | -0.009 | * |

The effect of independent variables on mediator variables, modeled as a linear mixed effects regression using Bayesian statistics. All mediator variables are continuous and standardized to a mean of 0 and standard deviation of 1. Thus the effect estimate indicates the mean difference in the mediator variable (eg RR) when comparing the two contrasts (e.g. two strains), expressed in units of standard deviation. The 95% confidence intervals indicate the lowest and highest esimates for the effect size. If the confidence interval crosses 0, then the effect is not significant.

Table S2: Dependence of network detectability on independent variables

Path C of the mediation analysis

| Independent Variable | Effect estimate | Confidence Interval (lower) | Confidence Interval (upper) | Significant? |
|---|---|---|---|---|
| strain (C3HeB/FeJ) | -0.06 | -0.20 | 0.09 | |
| sex (f) | 0.00 | -0.14 | 0.14 | |
| isoflurane (0.5% vs 0.23%) | -0.08 | -0.15 | 0.00 | * |
| isoflurane (1% vs 0.5%) | -0.23 | -0.30 | -0.15 | * |
| dexmedetomidine (0.05 vs 0.025) | -0.04 | -0.13 | 0.06 | |
| dexmedetomidine (0.1 vs 0.05) | -0.04 | -0.13 | 0.05 | |
| session (2 vs 1) | -0.02 | -0.07 | 0.03 | |
| session (3 vs 2) | 0.27 | 0.22 | 0.33 | * |
| Time after isoflurane change | -0.02 | -0.04 | 0.00 | * |
| strain : isoflurane (0.5% vs 0.23%) | -0.02 | -0.12 | 0.07 | |
| strain : isoflurane (1% vs 0.5%) | 0.02 | -0.06 | 0.10 | |
| sex : isoflurane (0.5% vs 0.23%) | -0.04 | -0.12 | 0.05 | |
| sex : isoflurane (1% vs 0.5%) | 0.01 | -0.07 | 0.09 | |
| strain : dexmedetomdine (0.05 vs 0.025) | -0.02 | -0.11 | 0.08 | |
| strain : dexmedetomidine (0.1 vs 0.05) | 0.20 | 0.10 | 0.30 | * |
| sex : dexmedetomidine (0.05 vs 0.025) | 0.22 | 0.13 | 0.31 | * |
| sex : dexmedetomidine (0.1 vs 0.05) | -0.21 | -0.30 | -0.12 | * |

The total effects of independent variables on network detectability, modeled as a linear mixed effects regression using Bayesian statistics. Network detectability is a spatial correlation that was Fisher-Z and log transformed. Thus the effect estimate indicates the mean difference in network detectability when comparing the two contrasts (e.g. two strains), expressed in units of log correlations. The 95% confidence intervals indicate the lowest and highest esimates for the effect size. If the confidence interval crosses 0, then the effect is not significant.

Table S3: Dependence of network detectability on independent variables while controlling for mediator variables

Path C' of the mediation analysis

| Independent Variable | Effect estimate | Confidence Interval (lower) | Confidence Interval (upper) | Significant? |
|---|---|---|---|---|
| strain (C3HeB/FeJ) | -0.147 | -0.298 | -0.005 | * |
| sex (f) | 0.048 | -0.079 | 0.173 | |
| isoflurane (0.5% vs 0.23%) | -0.032 | -0.107 | 0.044 | |
| isoflurane (1% vs 0.5%) | -0.183 | -0.256 | -0.105 | * |
| dexmedetomidine (0.05 vs 0.025) | 0.000 | -0.102 | 0.100 | |
| dexmedetomidine (0.1 vs 0.05) | -0.061 | -0.171 | 0.050 | |
| session (2 vs 1) | -0.039 | -0.101 | 0.023 | |
| session (3 vs 2) | 0.201 | 0.137 | 0.267 | * |
| Time after isoflurane change | -0.020 | -0.037 | -0.003 | * |
| strain : isoflurane (0.5% vs 0.23%) | -0.065 | -0.154 | 0.020 | |
| strain : isoflurane (1% vs 0.5%) | -0.015 | -0.095 | 0.068 | |
| sex : isoflurane (0.5% vs 0.23%) | -0.057 | -0.139 | 0.028 | |
| sex : isoflurane (1% vs 0.5%) | 0.005 | -0.076 | 0.083 | |
| strain : dexmedetomdine (0.05 vs 0.025) | -0.060 | -0.166 | 0.045 | |
| strain : dexmedetomidine (0.1 vs 0.05) | 0.294 | 0.172 | 0.411 | * |
| sex : dexmedetomidine (0.05 vs 0.025) | 0.273 | 0.166 | 0.383 | * |
| sex : dexmedetomidine (0.1 vs 0.05) | -0.331 | -0.439 | -0.225 | * |

The direct effects of independent variables on network detectability obtained when controlling for all mediator variables, modeled as a linear mixed effects regression using Bayesian statistics. Network detectability is a spatial correlation that was Fisher-Z and log transformed. Thus the effect estimate indicates the mean difference in network detectability when comparing the two contrasts (e.g. two strains), expressed in units of log correlations. The 95% confidence intervals indicate the lowest and highest esimates for the effect size. If the confidence interval crosses 0, then the effect is not significant.

Table S4: Mediator (instantaneous) variables that are predictive of network detectability above and beyond the independent variables.

Path B of the mediation analysis

| Mediator Variable | Effect estimate | Confidence Interval (lower) | Confidence Interval (upper) | Significant? |
|---|---|---|---|---|
| RR | 0.04 | 0.00 | 0.09 | * |
| RRV | 0.02 | 0.00 | 0.04 | * |
| RV | 0.02 | -0.02 | 0.06 | |
| HR | -0.03 | -0.08 | 0.02 | |
| PVI | -0.04 | -0.08 | 0.00 | |
| HRV | 0.06 | 0.01 | 0.11 | * |
| SpO2 | 0.05 | 0.02 | 0.08 | * |
| MeanFD | -0.04 | -0.07 | -0.01 | * |

The effects of mediator variables on network detectability when controlling for independent variables, modeled as a linear mixed effects regression using Bayesian statistics. Network detectability is a spatial correlation that was Fisher-Z and log transformed. The 95% confidence intervals indicate the lowest and highest esimates for the effect size. If the confidence interval crosses 0, then the effect is not significant.

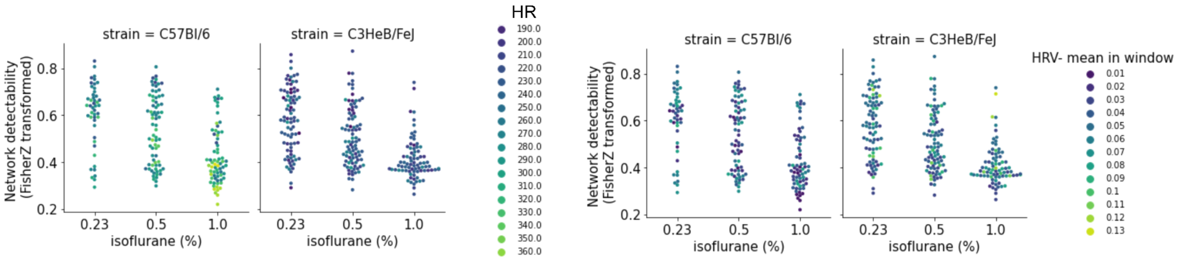


*Figure S4. The interaction between isoflurane dose and demographics on network detectability (y-axis) and physiological metrics (indicated via the point colors). Only the physiological metrics for which the interaction was significant are shown.*

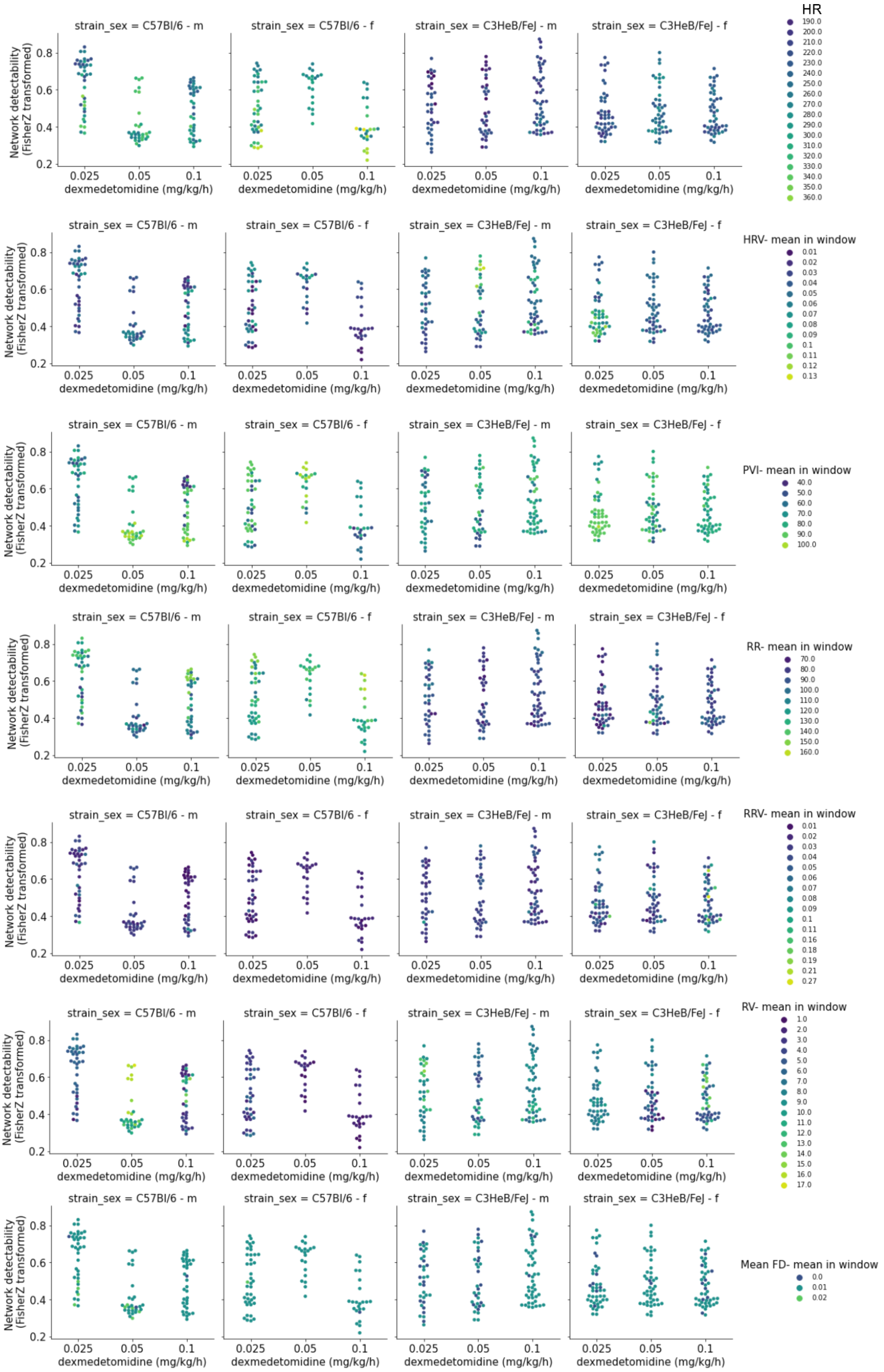
strain_sex = C57Bl/6 - m
strain_sex = C57Bl/6 - f
strain_sex = C3HeB/FeJ - m
strain_sex = C3HeB/FeJ - f
Network detectability
(FisherZ transformed)
0.2
0.4
0.6
0.8
0.025
0.05
0.1
dexmedetomidine (mg/kg/h)
HR
190.0
200.0
210.0
220.0
230.0
240.0
250.0
260.0
270.0
280.0
290.0
300.0
310.0
320.0
330.0
340.0
350.0
360.0
HRV- mean in window
0.01
0.02
0.03
0.04
0.05
0.06
0.07
0.08
0.09
0.1
0.11
0.12
0.13
PVI- mean in window
40.0
50.0
60.0
70.0
80.0
90.0
100.0
RR- mean in window
70.0
80.0
90.0
100.0
110.0
120.0
130.0
140.0
150.0
160.0
RRV- mean in window
0.01
0.02
0.03
0.04
0.05
0.06
0.07
0.08
0.09
0.1
0.11
0.16
0.18
0.19
0.21
0.27
RV- mean in window
1.0
2.0
3.0
4.0
5.0
6.0
7.0
8.0
9.0
10.0
11.0
12.0
13.0
14.0
15.0
16.0
17.0
Mean FD- mean in window
0.0
0.01
0.02

*Figure S5. The interaction between dexmedetomidine dose and demographics on network detectability (y-axis) and physiological metrics (indicated via the point colors). Only the physiological metrics for which the interaction was significant are shown.*

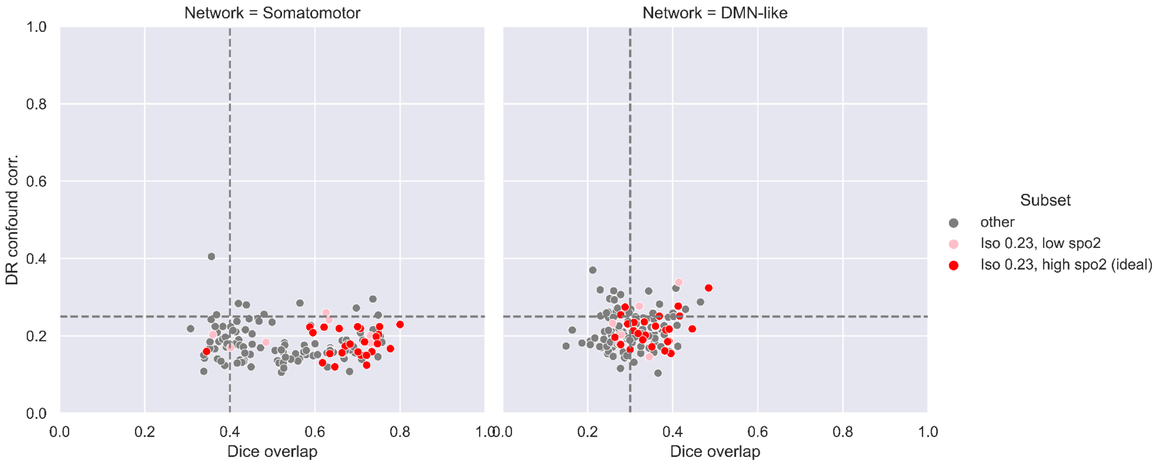


*Figure S6. The success rate of the recommended approach from the decision tree on the level of 8-minute windows. The Dice overlap and correlation with confounds were computed with RABIES after splitting each scan into 8-minute windows corresponding to each isoflurane dose. The dotted lines indicate the recommended thresholds, successful scans should have a dice overlap above 0.4 (or 0.3 for the DMN as it covers a smaller surface) and a correlation with confounds below 0.25 (Desrosiers-Grégoire et al. 2024). The points that were acquired under the recommended conditions , namely at 0.23% isoflurane and when the SpO2 was -0.7 standard deviations above the strain-specific average (>73% for C57Bl/6 and >86% for C3FeB/HeJ), are coloured in red. The pink points show the 8-min windows that were also acquired under 0.23% isoflurane but during which the SpO2 was below the threshold. Finally, all other windows at higher isoflurane levels are in gray. Within the recommended conditions, 96% of the windows contain a clear somatomotor network and 73% contain a clear DMN-like network (possibly the two scans lost due to a high correlation with confounds can be recovered following optimized confound correction).*
